# Antagonistic action of Siah2 and Pard3/JamC to promote germinal zone exit of differentiated cerebellar granule neurons by modulating Ntn1 signaling via Dcc

**DOI:** 10.1101/2022.06.21.497057

**Authors:** Christophe Laumonnerie, Maleelo Shamambo, Daniel R. Stabley, Tommy L. Lewis, Niraj Trivedi, Danielle Howell, David J Solecki

**Affiliations:** Department of Developmental Neurobiology, St. Jude Children’s Research Hospital, 262 Danny Thomas Place, Memphis, TN 38104; Aging & Metabolism Program, Oklahoma Medical Research Foundation, 825 NE 13th Street, Oklahoma City, OK 73104

**Keywords:** Germinal Zone, Polarity, Netrin1, Dcc, Pard3, JamC, Coincidence Detection

## Abstract

Germinal zone (GZ) exit is at the top of a cascade of events promoting the maturation of neurons and their assembly into neuronal circuits. Developing neurons and their progenitors must interpret varied niche signals like morphogens, guidance molecules, extracellular matrix, or adhesive cues to navigate GZ occupancy. How newborn neurons integrate multiple cell-extrinsic niche cues with their cell-intrinsic machinery in exiting a GZ is unknown. We establish cooperation between cell polarity-regulated adhesion and Netrin-1 signaling comprises a coincidence detection circuit repelling maturing neurons from their GZ. In this circuit, the Partitioning defective 3 (Pard3) polarity protein and Junctional adhesion molecule-C (JAM-C) adhesion protein promote, while the Seven in absentia 2 (Siah2) ubiquitin ligase inhibits, Deleted in colorectal cancer (DCC) receptor surface recruitment to gate differentiation linked repulsion to GZ Netrin-1. These results demonstrate cell polarity as a central integrator of adhesive- and guidance cues cooperating to spur GZ exit.

**Highlights:** -Netrin1 expressed in the cerebellar GZ ranges from an attractive to repulsive cue depending on differentiation status or extracellular matrix context.

-Modulation of DCC expression levels impacts the nature of the Netrin1 response and GZ occupancy status.

-Siah2, Pard3, and JamC function to modulate responsiveness to purified Netrin1 and cooperate with Dcc to regulate GZ exit.

-Pard3 and JamC promote both basal and Netrin1 stimulated DCC surface recruitment to control GZ exit and Netrin1 repulsion

## Introduction

During nervous system development, sequential events ensure the formation of the neuronal circuitry required for proper cognitive function. At the cellular level, maturing neurons navigate multifaceted tissue environments by interpreting cell-extrinsic factors that cooperate with cell-intrinsic programs to bring about the stereotypical progression of cells from neuronal progenitors to mature neurons with proper identity, location, and connectivity (Hatten, 1999). Flawed execution of these genetically encoded programs has drastic consequences for brain morphogenesis and circuit function, and it ultimately leads to a wide range of pediatric neurological disorders (Hakanen et al., 2019). There is compelling evidence that extrinsic factors, such as guidance molecules, and intrinsic programs, such as those involving neuronal polarity signaling or cytoskeletal organization, are important for germinal zone (GZ) exit, migration, and synaptogenesis in circuit formation. However, understanding how cell biological programs such as neuronal polarity signaling coordinate with extrinsic guidance signals remains a significant challenge as a result of a reductionist focus on how molecular players function in isolation and a lack of cell biological or imaging approaches by which to tackle this issue mechanistically. This study addressed this challenge by revealing how cell polarity signaling, cell adhesion, and the sensing of guidance molecules work in unison at the cell biological level to control when neuronal progenitor cells or newly differentiated neurons leave their germinal niche.

Granule neurons from the mouse cerebellar cortex have long been a system of choice for studying the process whereby neuronal progenitor replication in proliferative niches transitions to differentiation of neurons that migrate out of their GZ to a final laminar position (Chedotal, 2010; Hatten and Roussel, 2011; Leto et al., 2015). In the outer external granule layer (oEGL), granule neuron progenitors (GNPs) exhibit a low-polarity morphology. As they differentiate, newborn cerebellar granule neurons (CGNs) progressively extend a leading process parallel to the surface of the cerebellar cortex and start migrating along that axis in the inner external granule layer (iEGL). Eventually, CGNs extend a perpendicular process and the soma starts migrating radially on the surface of Bergmann glia fibers to reach the inner granule layer (IGL), leaving the axon behind in the molecular layer. This model is ideal for understanding how disparate cell biological processes act in unison. Recent work has shown that transcriptional and post-transcriptional regulation of polarity complex protein availability is critical for GZ exit. GNPs express the E3 ubiquitin ligase Siah2, which targets Pard3 for degradation (Famulski et al., 2010). Additionally, hypoxic pressure in the GZ and expression of the transcription factor Zeb1 transcriptionally repress Pard6α expression, thus maintaining cells in the EGL (Kullmann et al., 2020; Singh et al., 2016). As CGNs differentiate, polarity proteins are expressed, enabling one neurite to specialize as a leading process (Butts et al., 2014; Laumonnerie and Solecki, 2018) and promote the recruitment of junctional adhesion molecule type C (JamC) to the cell surface, thereby facilitating adhesion to the radial glial fibers and, thus, providing a substrate for migration through the molecular layer (Famulski *et al*., 2010). Despite the well-developed role of this polarity-dependent adhesion system in GZ exit, we still do not understand how this critical pathway is linked to classical guidance molecules that could functionally align cellular asymmetry to external reference points as newborn neurons take up their positions in the cerebellar circuit.

A large array of extrinsic cues have been identified as making short-range and long-range contributions to the cell cycle regulation of GNPs and the migration of CGNs (Leto *et al*., 2015). This myriad of signals raises the question of how a given neuron can filter out only those cues relevant to its behavior at a given time. We hypothesized that cellular polarization and polarity complex proteins could modulate the strength of extrinsic guidance signals by regulating the access of transmembrane receptors to the membrane surface. To test this idea, we investigated the function of the guidance molecule netrin-1 (Ntn1) in guiding CGNs out of the GZ. Ntn1 signaling is transduced by different transmembrane receptors of the immunoglobulin superfamily, like the deleted in colorectal cancer (Dcc) and Uncoordinated protein 5 (Unc5) transmembrane proteins, and is involved in axonal guidance and neuronal migration (Ackerman et al., 1997; Moore et al., 2007; Przyborski et al., 1998; Purohit et al., 2012). Based on the combination of receptors expressed by the cell, the signal can mediate attraction or repulsion behavior (Moore *et al*., 2007). Historically, in the cerebellar system, although Ntn1 is not involved in guiding the migration of GNPs from the upper rhombic lip, postnatal explants of the cerebellar cortex showed that parallel fiber extensions were repelled from a source of Ntn-1 (Alcantara et al., 2000). In addition, the mutant model for another Ntn1 receptor, Unc5c, has been described as possessing a smaller cerebellum and folia, ectopic cerebellar cells in the midbrain, and abnormal postnatal cerebellar migration (Ackerman *et al*., 1997).

In this study, we used the GZ exit of differentiating CGNs in the cerebellum as a model to study how extracellular guidance cue–receptor pairs, such as Dcc and Ntn1, a cellular polarity complex, and an adhesion protein can intersect, creating a coincidence detection circuit that modulates cellular migration. Our results reveal a localized source of Ntn1 in the EGL, where GNPs reside, and Unc5c upregulation promotes GZ exit, specifically of newly differentiated CGNs. Levels of Dcc receptors were found to be critical for tuning the sensitivity of the response. Using FACS-sorted populations of GNPs and CGNs, we show that differentiated neurons migrate away from a source of Ntn1, whereas progenitors show no preferential migration orientation. These observations led us to investigate how the access of transmembrane receptors to the membrane surface was regulated. Like Pard3, Dcc has been previously reported to be targeted for degradation by the proteasome by Siah proteins (Hu and Fearon, 1999; Hu et al., 1997; Li et al., 2015). Here we show that Siah2 promotes GZ occupancy by decreasing the amount of Dcc receptors at the membrane. Using super-resolution microscopy, we provide evidence that Dcc receptors reside near Pard3 and JamC at the membrane surface; additionally, we show that JamC, Pard3, and Dcc likely form a protein complex. We demonstrate that Pard3 and JamC functions are essential to a proper Ntn1-mediated migration phenotype in an ex vivo assay.

Necessity–sufficiency testing revealed that Pard3 promotes Dcc receptor clustering in response to netrin-1, whereas JamC controls the basal levels of Dcc at the surface. Thus, Pard3 and JamC regulate the sensitivity of newly differentiated CGNs to the netrin gradient by adjusting the number of Dcc receptors recruited to the cell surface and cytoskeleton. This new paradigm for cooperativity between disparate cell biological processes illustrates that coincidence detection between JamC and polarity-dependent adhesion and sensitivity to a repulsive netrin-1 work in tandem to facilitate GZ exit, with the polarity pathway concentrating Dcc at the cell surface by stabilizing and/or promoting transmembrane receptor exocytosis.

## Results

Nnt1 signaling has been implicated in guiding early aspects of cerebellar morphogenesis, such as repulsion of CGN axons and the positioning of the boundary of the forming EGL (Ackerman *et al*., 1997; Alcantara *et al*., 2000), making it an excellent guidance molecule model for assessing cooperativity with polarity proteins. We first analyzed the expression of Ntn1 and its receptors Dcc and Unc5c at the protein level in postnatal day 7 (P7) mice. Immunohistochemical staining revealed that Ntn1 is present in the oEGL complementary to Cntn2 CGN marker expression in the iEGL. In contrast, when the Atoh1::eGFP marker was used to highlight GNPs in the oEGL, immunohistochemical staining for Ntn1 receptors showed that Dcc and Unc5c are expressed in the iEGL. Whereas Dcc expression is maintained in the IGL, Unc5c appears to be more restricted to the iEGL (Figure 1A). These expression patterns suggest that Ntn1 and its receptors are well-positioned to affect CGN GZ exit and/or occupancy.

**Figure 1.**
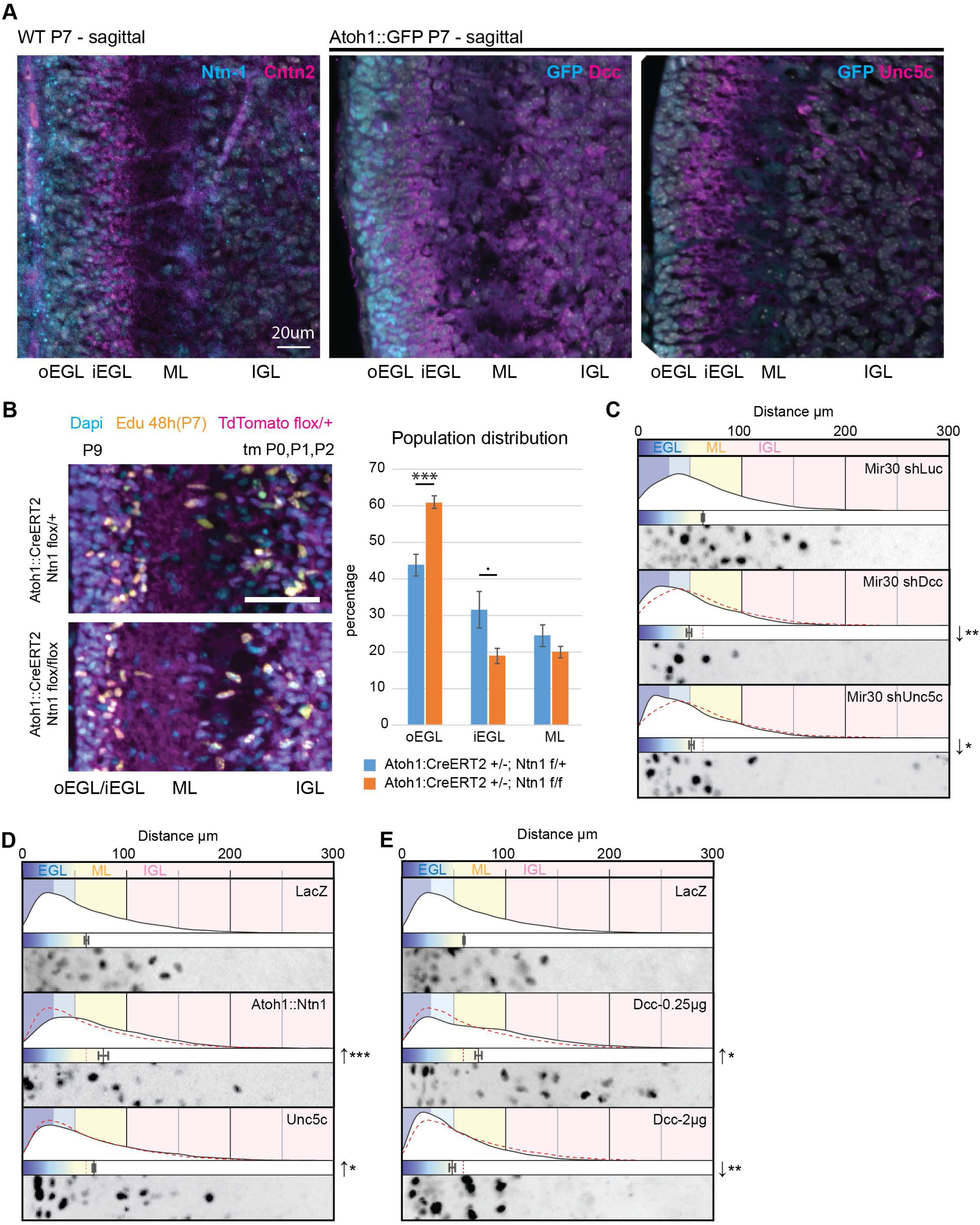
(A) Immunohistochemical staining of sagittal cryo-sections of wild-type and Atoh1-GFP mouse cerebella at P7. From left to right, the sections were stained used antibodies against Ntn1 (cyan) and Cntn2 (magenta), GFP (cyan) and Dcc (magenta), and GFP (cyan) and Unc5c (magenta), respectively, and were counterstained with Dapi (white). (B) Sagittal cryo-sections through the cerebella of P9 Rosa::TdTomato^fl/+^; Atoh1::CreERT2^+/-^; Ntn1^flox/wt^ or Ntn1^flox/flox^ animals after three doses of tamoxifen (tm) at P0, P1, and P2 and a single injection of Edu at P7. Sections were stained for Edu, and the radial distance from the edge of the Edu^+^ nucleus was measured and plotted. oEGL: 0–30 µm; iEGL: 30–50 µm; ML: 50–100 µm. (C)–(E). Results of ex vivo slice culture assays under different conditions. In each case, the top curve shows the entire distribution of the radial distances of H2B-positive electroporated nuclei from the edge of the slice in replicates. Below this is a plot of the average radial distance from the edge among replicates, and below this is a micrograph representative of the nuclear distribution in that assay. All are displayed on the same scale, representing a distance from 0 to 300 µm. In addition to H2B-Cherry, the following constructs were electroporated: in (C), Mir30 shLuc (control), Mir30 shDcc, and Mir30 shUnc5c; in (D), LacZ (control), Atoh1::Ntn1, and Unc5c; in (E), LacZ (control), Dcc at 0.25 µg, and Dcc at 2 µg. Each respective control is represented by a dashed red line in the distribution plot. Scale bars in (A) and (C) represent 20 µm and 10 µm, respectively. Abbreviations: oEGL, outer external granule layer; iEGL, inner external granule layer; EGL, external granule layer; ML, molecular layer; IGL, internal granule layer. In (B) through (E), the error bars represent the SEM. Statistics: *p ≤ 0.05; **p ≤ 0.01, ***p ≤ 0.005, as assessed by a Student *t*-test in (C) and an ANOVA followed by a Dunnett post hoc test in (B), (D), and (E) against the respective controls. See also Table S1.

We next assessed whether Ntn1 function affected GZ exit in the developing cerebellum in vivo by using a Cre-inducible system to generate Atoh1::CreERT2/Ntn1^lox/lox^; Rosa::TdTomato^lox/wt^ mice. In vivo GZ pulse-chase analysis was conducted in animals to whom tamoxifen was administered daily from P0 to P3, using one dose of 5-ethynyl-2’-deoxyuridine (EdU) injected at P7. The EdU GZ pulse labels the proliferative GNPs of the oEGLin the cerebellar cortex when injected at P7. The position of Edu/TdTomato-positive cells in tissues collected at later stages of development can be used as a proxy to analyze the GZ exit and migration of the CGN progeny of GNPs in vivo. At P9, the average distance from the cerebellar surface of Edu/TdTomato-positive nuclei, which is a measure of GZ exit and subsequent migration, was lower in Atoh1::CreERT2/Ntn1^lox/lox^; Rosa::TdTomato^lox/wt^ mice than in their Atoh1::CreERT2/Ntn1^lox/WT^; Rosa::TdTomato^lox/wt^ littermates, showing that deleting Ntn1 from the EGL results in the inhibition of GZ exit (Figure 1B).

Given the expression of Dcc and Unc5c in differentiated CGNs, we next assessed the function of these receptors in GZ exit by using the ex vivo slice culture assay developed in our laboratory. Silencing of both Unc5c and Dcc via Mir30 shRNA expression significantly reduced the average migration distance from the pial surface after 48 h in culture and increased the proportion of cells that remained in the EGL (Figure 1C). These results are consistent with studies showing that Dcc/Unc5 heterodimers mediate chemorepulsion from a source of Nnt1 (Moore *et al*., 2007), buttressing our finding that Ntn1 loss of function led to enhanced GZ occupancy. Interestingly, the average migration speed was not affected by Dcc and Unc5c silencing in dissociated CGN cultures (Figure S1A), showing that the loss of GZ exit in silenced cells is not due to defective cell motility.

Given the necessity of Ntn1, Dcc, and Unc5c for CGNs to take up positions outside the EGL, we next tested whether they were sufficient to induce GZ exit. Expressing Ntn1 in ex vivo slices by using the Atoh1 regulatory sequence to restrict its expression to the oEGL increased the average distance of electroporated nuclei from the pial surface and reduced EGL occupancy (Figure 1D). Elevated expression of Unc5c, which is expressed only as CGNs differentiate, resulted in a similar precocious GZ exit phenotype (Figure 1D). Expression of Dcc exhibited a concentration-specific phenotype: a mild elevation of Dcc expression promoted GZ exit, whereas at higher concentrations, cells remained in the GZ (Figure 1E). This phenotypic transition was gradual (Figure S1B) and, based on the literature, could reflect the fact that increasing the number of Dcc receptors at the cell surface favors Dcc homodimer formation and attraction to a source of Ntn1 (Moore *et al*., 2007; Xu et al., 2014), whereas when the number of Dcc receptors is low, they may cooperate with endogenous Unc5, promoting repulsion. These results show that Dcc expression and the regulation of its availability at the membrane surface appear to be critical to modulating the GZ exit response to Ntn1 in the oEGL.

To assess whether Ntn1 signaling directly affected CGN migration direction, we used an in vitro system to control the source of this guidance cue. Channel microslides can set and hold a gradient for several hours, enable different substrate coatings to be placed on the glass base of the slide, and are optically compatible with time-lapse microscopy. Dissociated cerebellar neurons were nucleofected and seeded into the channel of a microslide and incubated for 24 h. Ten minutes before the start of an imaging experiment, the medium in the channel was replaced, and that in the left half of the slide was replaced with medium containing 200 ng/mL of recombinant Ntn1 or its equivalent in volume in 1× PBS as a control. Fluorescein was also added to enable visualization of the gradient status over time (Figure 2A). The motion of H2b-mCherry–labeled nuclei was assayed via 2 h of live-cell time-lapse microscopy followed by computational tracking of nuclei to determine the cell response to Ntn1 (Figure 2B). For simplicity, we expressed the tracking results as variations from a 50%:50% distribution in the x-axis of the slide, as there were no specific alterations in the relative endpoint displacement in the gradient y axis. Although the endpoint distribution of migration on the x-axis showed no preferential direction in control cells, the unilateral addition of Ntn1 resulted in an attractive shift toward the source of Ntn1 (Figure 2C). A chi-square statistical test revealed that the pattern of attraction observed with unilateral addition of Ntn1 was significantly different from a 50%:50% distribution (Figure 2D). To validate the dependency of migration on Ntn1 signaling, we nucleofected constructs to modify the expression levels of the Ntn1 receptors. On laminin-coated slides, a gain of function of Dcc maintained the attraction phenotype observed in controls, whereas the gain of function of Unc5c reversed the phenotype to a repulsive one, with cells migrating away from the source of Ntn1 (Figure 2D). Mir30-based shRNA knockdown of both Dcc and Unc5c resulted in the loss of responsiveness to the source of Ntn1 (Figure 2D). Overall, these results confirm that the Ntn1 receptors Dcc and Unc5c are essential to reorienting the migration of dissociated CGNs in response to a source of Ntn1.

**Figure 2.**
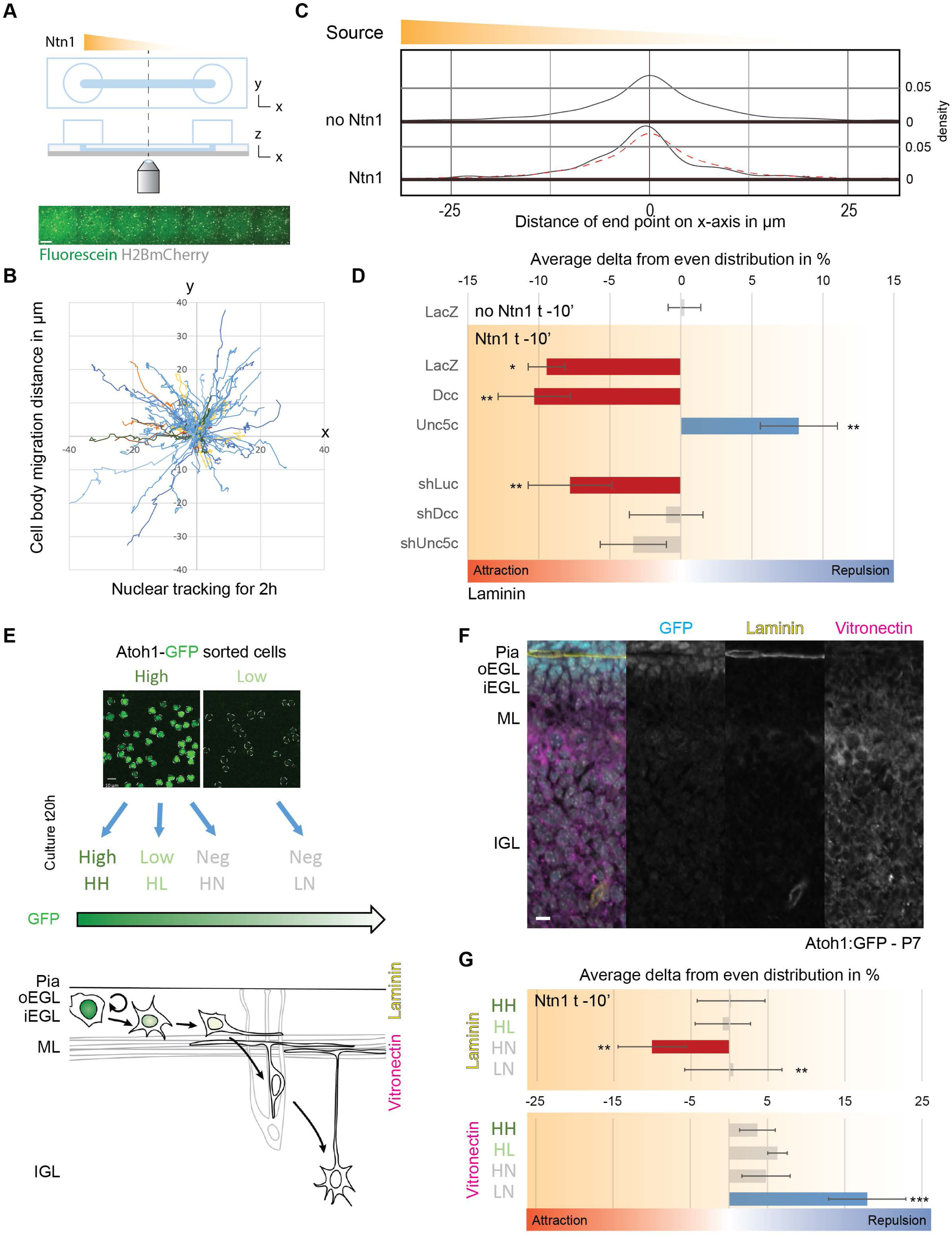
(A) Schematic representing the channel microslides used to assess migratory behavior in response to the unilateral addition of Ntn1 from time-lapse imaging. The panel below shows tiled pictures of the dissociated cerebellar granule neurons nucleofected with H2B-mCherry (white) within a fluorescein gradient (green) created in the middle area of the channel microslide. (B) Example of manual tracking of nuclear motion during the 2-h time-lapse imaging session, here without the addition of Ntn1. Each track is centered at t0, and motions are displayed as the relative displacement from the the origin. The x-axis is aligned with the channel length, as depicted in (A). (C) Graph representing the density distribution of the endpoint displacements on the x-axis across all of the tracked migration in the “no Ntn1” control (upper graph and red dashed line in lower graph) (n = 338) and with the unilateral addition of Ntn1 as depicted by the “source” bar at top (n = 379). The gradient was set 10 min before the start of the 2-h time-lapse imaging session. (D) Chart representing the average variation across replicates in the endpoint nuclear displacement on the x-axis from an even probability of 50%:50%, with negative values representing an attraction to the source of the gradient (no Ntn1 or Ntn1) (in red when statistically significant) and positive values representing repulsion (in blue when statistically significant). Dissociated cerebellar granule neurons were plated on laminin-coated channel microslides 24 h before tracking began, and the gradient was set 10 min before the start of image acquisition. Cells were nucleofected with H2B-mCherry, GPI-pHluorin, and the following: LacZ (control), Dcc, Unc5c, shLuc (control), shDcc, or shUnc5c. (E) Top. Plated cells after dissociation from cerebella of Atoh1-GFP mice and sorting by FACS into high-GFP-intensity and low-GFP-intensity populations, showing Atoh1-GFP signal (green). Bottom. Schematic representation of the cerebellar cortical layers of a P7 Atoh1-GFP mouse, showing the decrease in GFP intensity during the differentiation process and the migration of a GNP from the oEGL to the IGL and the corresponding switch of the extracellular matrix from laminin-rich to vitronectin-rich. (F) Immunohistochemical staining of a sagittal cryo-section of Atoh1-GFP mouse cerebellum at P7 with antibodies against GFP(cyan), laminin (yellow), and vitronectin (magenta). (G) Chart similar to that in (D) representing the bias in the nuclear migration of different populations of FACS-sorted Atoh1-GFP mouse cerebellar neurons plated on channel microslides coated with either laminin or vitronectin after unilateral addition of Ntn1 10 min before the start of the 2-h time-lapse imaging. Scale bars in (A), (E), and (F) represent 100, 10, and 10 µm respectively. Abbreviations: oEGL, outer external granule layer; iEGL, inner external granule layer; EGL, external granule layer; ML, molecular layer; IGL, internal granule layer. In (D) and (G), the error bars represent the SEM. Statistics: *p ≤ 0.05, **p ≤ 0.01, ***p ≤ 0.005, as assessed by chi-square test against an even probability of 50%:50%. See also Table S1.

Given the differentiation-specific expression of Dcc and Unc5c, we were curious as to whether GNPs and CGNs at various differentiation stages had distinct responses to Ntn1. We took advantage of the knock-in mouse line in which EGFP is fused in-frame with the Atoh1 transcription factor that is expressed specifically in GNPs (Figure 2E). Atoh1-EGFP fluorescence had two uses in our experiment: 1) Atoh1-EGFP fluorescence could be used to sort dissociated EGL cells into GNPs (Atoh1-high, Pax6^+^) and CGNs (Atoh1-low, Pax6^+^) (Figure S2A), with Atoh1-negative cells from the initial sort being Pax2^+^ interneurons (Figure S2B); and 2) the intensity of Atoh1-EGFP in time-lapse imaging assays reported the differentiation status of the purified CGN lineage cells in real time. In the Ntn1 migration assay, Atoh1-high and Atoh1-low populations were plated on microslides 24 h before time-lapse imaging of their response. We also coated microslides with extracellular matrix components that are enriched near the oEGL (laminin) or iEGL (vitronectin) to better recapitulate conditions near the EGL niche (Figure 2F). Our time-lapse imaging revealed that the Atoh1-high population maintained a high level of Atoh1-GFP (true GNPs; designated HH cells), transitioned to a low level of Atoh1-GFP (intermediate GNPS; designated HL cells), or lost Atoh1-GFP altogether (recently differentiated CGNs; designated HN cells) (Figure 2E). Cells coming from the “low” sorted population remained negative for Atoh1-GFP and represented more mature CGNs (designated LN cells in Figure 2E). On both substrates, the HH and HL populations of Atoh1-GFP–positive cells showed no significant directional bias when migrating in the Ntn1 gradient (Figure 2G). On laminin, the HN population of newly differentiated CGNs migrated significantly toward the source of Ntn1 but showed no preference on vitronectin (Figure 2G). Conversely, the LN population of more mature CGNs migrated away from the Ntn1 source on vitronectin, which mimics the extracellular matrix components in the iEGL and the molecular layer. The behavior of the LN population on laminin was heterogeneous. A comparison to an even distribution with a chi-square test (Figure 2G) revealed that some cells had an attraction response (Figure S2C, magenta arrowhead) whereas others had a strong repulsion response (Figure S2C, cyan arrowhead), indicating that HL cells exposed to laminin could have a slower transition from attraction to repulsion. It is worth noting that vitronectin promotes CGN differentiation (Ong et al., 2020; Pons et al., 2001), whereas laminin, in some contexts, maintains GNPs and CGNs in an immature state (Ong and Solecki, 2017; Ong *et al*., 2020). Taken together, our in vivo, ex vivo, and microslide experiments show that the level of Dcc protein is critical for the regulation of GZ exit by Ntn1 signaling (Figure 1E) and that GNPs and CGNs display distinct responses to the presence of Ntn1 in the EGL GZ that differ according to their maturation status (Figure 2G).

We were intrigued by the differentiation-specific expression of Dcc in CGNs and by the fact that Dcc expression levels modulated the response to Ntn1 in the ex vivo cerebellar slice assay. Accordingly, we assessed how Dcc levels were controlled during CGN differentiation. Previous work from our laboratory has shown that Siah2 (seven in absentia homolog 2), a Ring-domain E3 ubiquitin ligase, is strongly expressed in GNPs and that it regulates GZ occupancy by targeting for degradation key proteins of the polarity complex, primary cilia, and hypoxia pathway (Famulski *et al*., 2010; Kullmann *et al*., 2020; Ong and Solecki, 2017; Ong *et al*., 2020). Furthermore, we show here that Siah2 expression decreases with Atoh1-GFP fluorescent signal as CGNs differentiate and start to migrate deeper in the cerebellum (Figure S3A). Indeed, the sorted cells we used for our channel microslide experiments (e.g., HH, HL, HN, and LN cells defined in the last section) significantly vary in their relative Siah2 expression level in addition to their behavior in a Ntn1 gradient (Figure 3A). The Dcc protein carries the canonical degron motif Pro-X-Ala-X-Val-X-Pro, which is recognized by Siah proteins within its intracellular P2 domain (House et al., 2003) (Figure 3B). This motif is well conserved in organisms ranging from *Xenopus* to humans (Figure S3B) and has been shown to promote specific proteasomal degradation of Dcc by Siah proteins (Hu *et al*., 1997). To confirm the specificity of Siah2-mediated degradation of Dcc, we co-expressed Siah2 with Dcc fused to pHluorin (Dcc-pH) in HEK293T cells. We also co-expressed Siah2 with a mutant form of Dcc-pH, Dcc-pH NXN, in which the last valine and proline of the Siah degron motif are replaced by two arginines, thereby abrogating Siah degradation (House *et al*., 2003) (Figure 3C). Whereas Siah2 decreased the level of Dcc-pH protein, that of Dcc-pH NXN remained unaffected by co-expression with Siah2. Furthermore, immunoprecipitation with an antibody against GFP showed that the K48-linked ubiquitin signal was more abundant when Dcc-pH was co-expressed with Siah2 but was absent when Dcc-pH NXN was co-expressed with Siah2 (Figure 3D). Therefore, Siah2 degradation of Dcc is a likely mechanism for restricting Dcc expression to CGNs, as these cells lack Siah2 expression.

**Figure 3.**
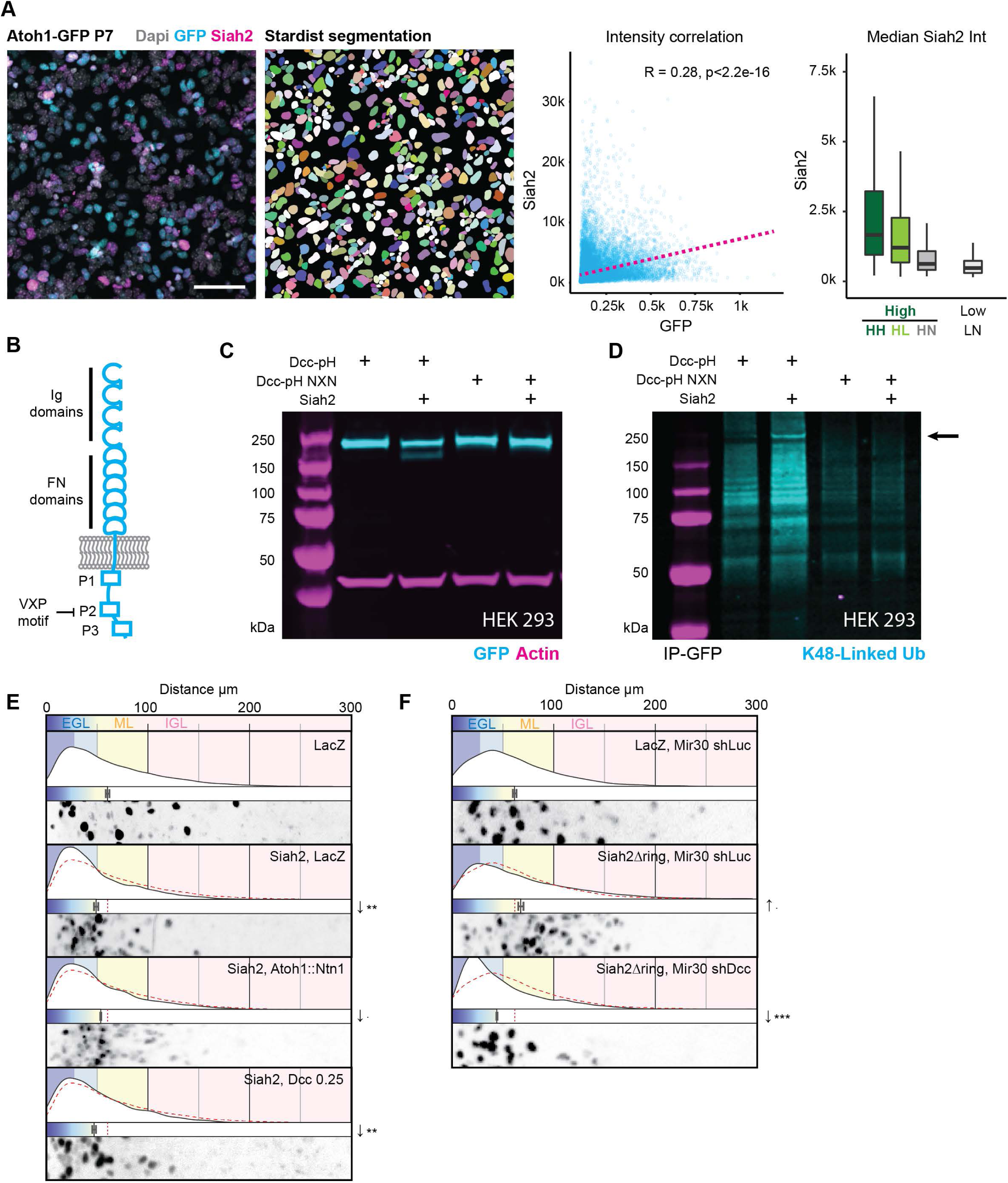
(A) (Left) Immunocytochemistry staining of the “High” and “Low” (not shown) population of Atoh1-GFP FACS-sorted CGNs from P7 mouse cerebellum and cultured for 24h, against GFP (Cyan) and Siah2 (Magenta), counterstained with Dapi (Gray). Picture on the right show an example of color-coded object after Stardist segmentation of the nucleus. First Graph shows the significative Pearson correlation between GFP and Siah2 signal intensity in all segmented cell bodies (“High” and “Low”). Boxplot show distribution of Siah2 intensity for the 4 populations previously defined in Figure 2 of granule neurons based on GFP intensity from the “High” sort: HH (->“High”), HL (->“Low”) and HN (->“Neg” (negative)) and the “Low” sort: LN (->”Neg”). (B) Schematic representing the different domains of Dcc, a single-pass transmembrane receptor for Ntn1 with four Immunoglobulin-like (Ig) domains and 6 fibronectin-like (FN) domains on its extracellular side and three intracellular domains, P1, P2, and P3. The schematic indicates the location of the canonical Siah2 degron motif VXP within P2. (C) Western blot of lysates of HEK293T cells after lipofection with Dcc-pHluorin (Dcc-pH), Dcc-pHluorin NXN mutant (Dcc-pH NXN), and Siah2. The blot was immunostained with antibodies against GFP (cyan) and actin (magenta). (D) Western blot of HEK 293 cells lysates immunoprecipitated with an antibody against GFP after the cells were lipofected with Dcc-pHluorin (Dcc-pH), Dcc-pHluorin NXN mutant (Dcc-pH NXN), and Siah2. Cells were treated with MG132 for 6 h before lysis. The blot was immunostained with an antibody against K48 ubiquitin (cyan). The arrow points to the lack of K48-ubiquitin–positive staining with the Dcc-pH NXN construct. (E) and(F). Results of ex vivo slice culture assays under different conditions. In each case, the top curve shows the entire distribution of the radial distances of H2B-positive electroporated nuclei from the edge of the slice in replicates. Below this is a plot of the average radial distance from the edge among replicates, and below this is a micrograph representative of the nuclear distribution in that assay. All are displayed on the same scale, representing a distance from 0 to 300 µm. In addition to H2B-Cherry, the following constructs were electroporated: in (E), LacZ (control), Siah2 and LacZ, Siah2 and Atoh1::Ntn1, and Siah2 and Dcc at 0.25µg; in (F), LacZ and Mir30 shLuc (control), Siah2Δring and Mir30 shLuc, and Siah2Δring and Mir30 shDcc. Scale bar in (A) represents 50 µm. Abbreviations: EGL, external granule layer; ML, molecular layer; IGL, internal granule layer. In (E) and (F), the error bars represent the SEM. Statistics: p ≤ 0.1, *p ≤ 0.05, **p ≤ 0.01, ***p ≤ 0.005, as assessed by an ANOVA followed by a Dunnett post hoc test against the respective controls. See also Table S1.

We next assessed whether Siah2 regulation of Dcc protein levels was relevant to CGN GZ exit by conducting epistasis analysis in our ex vivo cerebellar slice and microslide assays. Elevated Siah2 expression inhibited enhanced GZ exit that was elicited by Ntn1 and Dcc gain of function, showing that Siah2 antagonizes Dcc function in GZ exit (Figure 3E). Indeed, Dcc silencing inhibited the boost in GZ exit that is elicited by overexpression of a dominant-negative form of Siah2 lacking the Ring domain (Siah2ΔRing) (Famulski *et al*., 2010; Hu and Fearon, 1999), showing that the additional GZ exit that accompanies Siah2 inhibition relies on Dcc function, as would be expected from the Siah2–Dcc antagonism (Figure 3F). Taken together, these results suggest that Siah2 expression in GNPs actively hampers GZ exit by targeting Dcc for degradation.

Our previous work showed that Siah2 regulates GZ exit by acting as a negative regulator of partitioning defective (Pard) cell polarity pathways that promote GZ exit. Given that Dcc has been described as a neuronal polarity inducer (Adler et al., 2006) that, according to our findings, regulates GZ exit and is also negatively regulated by Siah2, we were curious as to whether there was a relation between the Pard complex and Dcc. Partition deficient protein 3 (Pard3) also contains the Siah2 degron motif, and the proteasomal degradation of the latter prevents Pard3 from recruiting JamC to the cell membrane surface, with the result that CGNs accumulate in the GZ (Famulski *et al*., 2010). Immunocytochemical staining of dissociated CGNs plated on laminin revealed areas in which the Dcc signal overlapped with the signal for Pard3 or JamC or both (Figure S4B). We designed fusion proteins carrying reporters to enable us to image all three proteins in live neurons with super-resolution. CGNs expressing Dcc-pHluorin, JamC-SNAP (stained with LAMPshade magenta, (Brown et al., 2021)), and Halo-Pard3 were imaged with an Airyscan microscope, which has approximately 100-nm XY resolution. The pHluorin fluorescent protein fused to Dcc and the LAMPshade magenta dye that stained JamC-SNAP share a pH-dependent fluorescence such that we imaged only the population of Dcc and JamC that was exocytosed to the extracellular face of the plasma membrane, which has a neutral pH when compared to the cell interior. When expressed together, the Dcc signal overlapped with that of JamC at the cell surface in locations where cells formed the point of contact with other cells in large, bright adhesion plaques (Figures 4A–4C; Figure S4C). Whereas the JamC signal appeared stable, Dcc was more dynamic in the middle of the JamC domain and coalesced at the edge of some JamC-labeled adhesions (Movie S1). The DCC signal sometimes coincided with the underlying Pard3 signal in fast-moving retrograde or stable Dcc clusters (Figures 4A–4C; Figure S4D; Movie S2). When Dcc, JamC, and Pard3 were co-expressed together, Pard3 accumulated below the large plaque of JamC that also overlapped with a brighter Dcc signal (Figure 4A, white arrowhead). It is important to note that Dcc did not localize exclusively to JamC/Pard3 structures and was distributed as a diffuse signal or in a clustered (brighter) structure at the membrane surface. Bright Dcc clusters also often accumulated around a JamC/Pard3 bright adhesion between two cells (Figure 4B, hollow arrowhead). After Ntn1 was added to the medium, Dcc aggregated in clusters (Figure 4C, cyan arrowhead), some of which were recruited to JamC/Pard3 plaques (Figure 4C, hollow arrowhead). To test whether Dcc interacted in a complex with JamC, we expressed these proteins in HEK293T cells and showed that JamC-Halo was present in the pull-down when Dcc-pHluorin was immunoprecipitated with an anti-GFP antibody. The JamC– Dcc interaction appeared to require the Dcc extracellular domain, as JamC was able to pull-down the extracellular domain of Dcc but not the intracellular domain alone (Figure S4D). Dcc-pHluorin could also pull down Pard3-Halo (Figure S4E), but we were less confident about this interaction because Pard3-Halo showed periodic binding to the immunoprecipitation beads. Regardless, the proximity of Dcc, JamC, and Pard3 in CGNs and the interactions detected in immunoprecipitation experiments suggest that Dcc forms a complex with polarity-related proteins such as JamC and Pard3.

**Figure 4.**
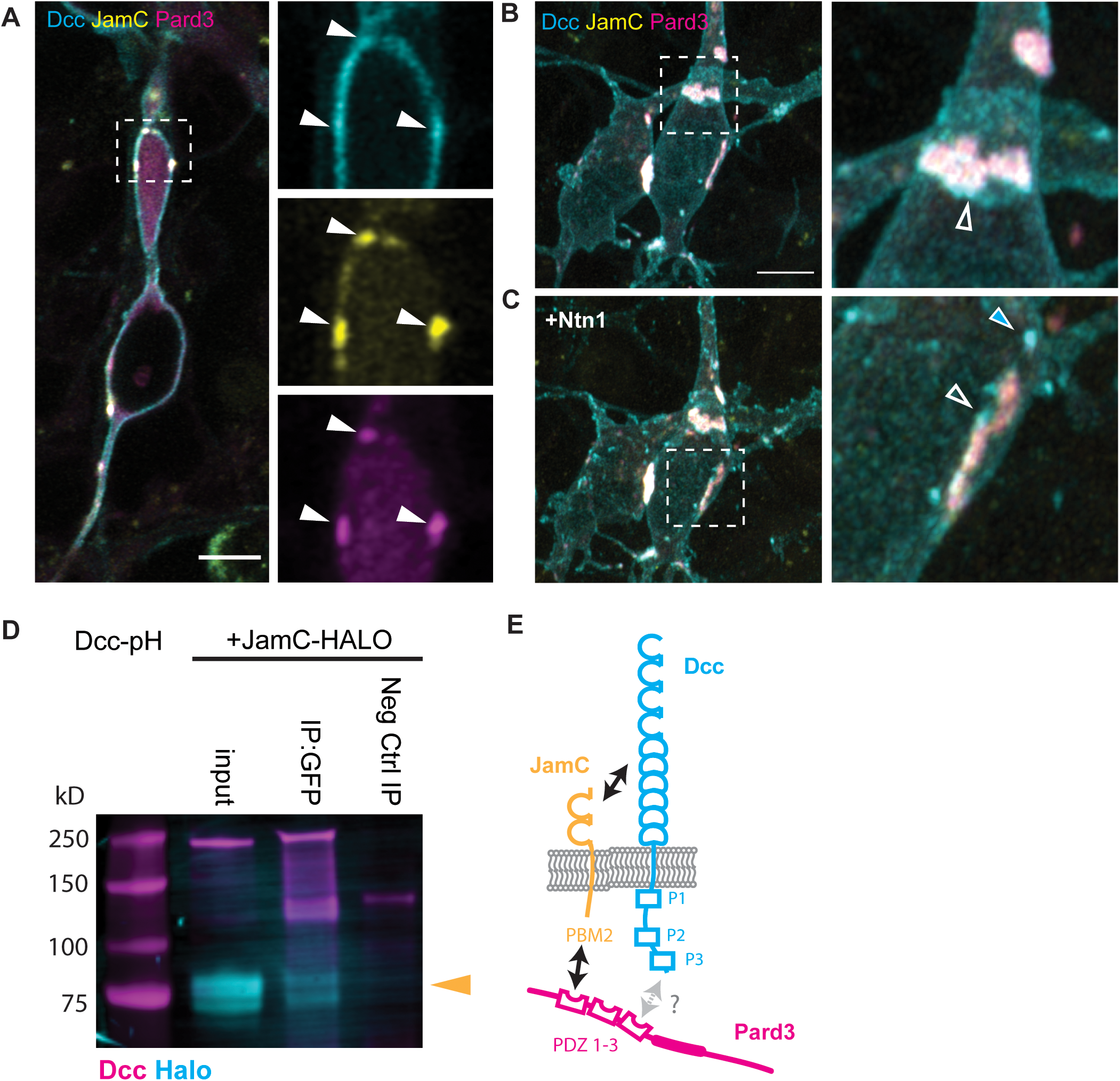
(A) –(C) Airyscan confocal live imaging of CGNs nucleofected with Dcc-pHluorin (cyan), JamC-SNAP (yellow), and Halo-Pard3 (magenta) constructs. Phluorin and the SNAP dye used here are both pH sensitive, emitting fluorescence only if exposed to a neutral pH. Thus, they show only proteins currently at the cell membrane surface. (A) A single focal plane 200 nm in thickness, showing overlap of the Dcc signal with the signals for JamC and Pard3 in the proximal dilation of a CGN (indicated by white arrowheads in the magnification). (B and C) Maximum projection of two CGNs forming an adhesion before (B) and after (C) the addition of Ntn1 to the medium. At the point of contact between the two cells, the Dcc signal overlapped the Pard3 and JamC signals (see magnifications). Additionally, Dcc accumulated at the edge of the adhesion without overlapping with Pard3 or JamC (arrowheads in B and C). Five minutes after the addition of Ntn1 at 200ng/L (C), the number of bright Dcc clusters (blue arrowhead) at the membrane surface increased and some newly formed clusters were recruited to the periphery of the JamC/Pard3/Dcc-positive adhesion (arrowhead in C). (D) Western blot of lysates of HEK293T cells after lipofection with Dcc-pHluorin (JamC-pH), and JamC-HALO. Cell lysates were immunoprecipitated with an antibody against GFP (IP: GFP) or a negative-control rabbit IgG (Neg Ctrl IP). The blots were immunostained with antibodies against Dcc and HALO-tag. Yellow arrowhead highlight the expected band size for JamC-HALO. (E) Schematic representing the predicted interactions between Dcc, JamC, and the polarity protein Pard3. Pard3 recruits JamC to the cell surface via its PDZ domain PDZ 1, which interacts with the Class 2 PDZ binding motif at the C-terminal of JamC. JamC interacts with the Dcc extracellular domain. Additionally, the PDZ 3 domain of Pard3 is predicted to interact with a Class 1 PDZ binding motif like the one present on the Dcc intracellular domain (X-S/T-X-φ-COOH). Scale bars in (A) and (B) represent 5 µm.

We next conducted epistasis experiments to test whether Pard3 and JamC influenced the effect of Ntn1 on GZ exit through Dcc in an ex vivo slice assay. Knockdown of both Pard3 and JamC was previously reported to prevent radial migration of CGNs and GZ exit (Famulski *et al*., 2010) (Figure 5A). Overexpression of Ntn1 in the GZ or mild expression of Dcc stimulated GZ exit in Pard3-silenced cells, meaning that the loss of Pard3 could be rescued by increasing the Ntn1 signaling (Figure 5A). In contrast, the phenotype observed with loss of JamC could not be rescued by overexpression of Ntn1, and mild overexpression of Dcc only restored GZ exit to the control level (Figure 5A). We then tested whether the knockdown phenotype of Dcc could be rescued by overexpression of Pard3 or JamC. Pard3 expression that promoted GZ exit also rescued knocked-down Dcc, restoring the level to that in the controls, and JamC expression rescued migration, restoring it to that seen in the control group (Figure 5B). In channel microslide assays, overexpression of both Pard3 and JamC in an unsorted population of dissociated CGNs plated on laminin led to a switch in the migratory response from attraction to repulsion with respect to a source of Ntn1 (Figure 5C), whereas knockdown left the cellular migration unbiased with respect to the Ntn1 source (Figure 5C). Taken together, these results suggest that Pard3 and JamC contribute to the way in which CGNs integrate Ntn1 signaling through Dcc, as their levels affect the outcome with respect to cellular migration and they are necessary for generating an oriented GZ repulsion migratory response (Figure 5C). This suggests that Pard3 and JamC synergize with Dcc to integrate the Ntn1 signal that guides CGNs out of the GZ as they differentiate.

**Figure 5.**
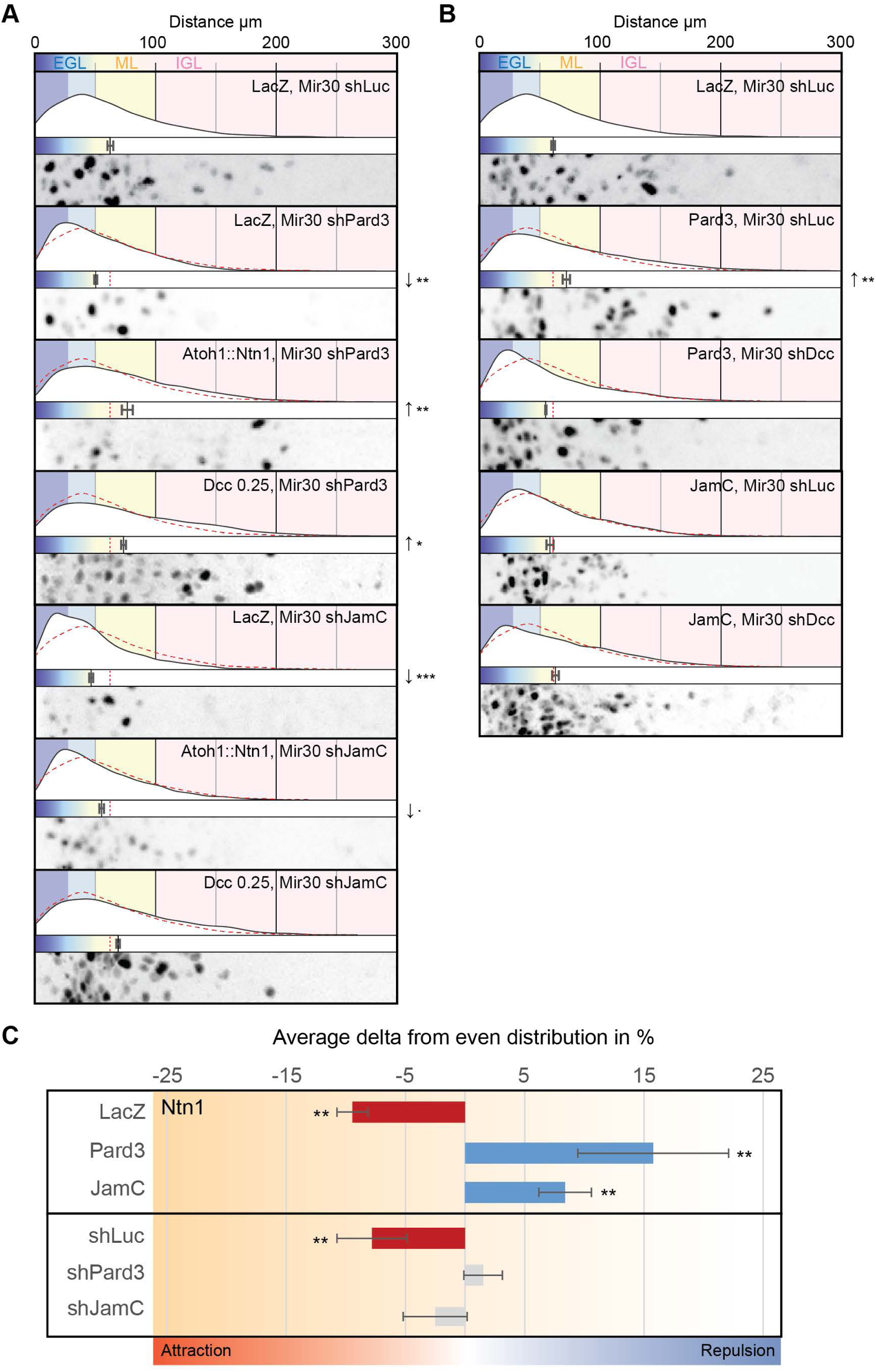
(A) and (B) Results of ex vivo slice culture assays under different conditions. In each case, the top curve shows the entire distribution of the radial distances of H2B-positive electroporated nuclei from the edge of the slice in replicates. Below this is a plot of the average radial distance from the edge among replicates, and below this is a micrograph representative of the nuclear distribution after 48 h in culture. All are displayed on the same scale, representing a distance from 0 to 300 µm. In addition to H2B-Cherry, the following constructs were electroporated: in (A), LacZ, Mir30 shLuc (control); LacZ, Mir30 shPard3; Atoh1::Ntn1, Mir30 shPard3; Dcc at 0.25 µg, Mir30 shPard3; LacZ, Mir30 shJamC; Atoh1::Ntn1, Mir30 shJamC; and Dcc at 0.25 µg, Mir30 shJamC; in (B), LacZ, Mir30 shLuc (control); Pard3, Mir30 shLuc; Pard3, Mir30 shDcc; JamC, Mir30 shLuc; and JamC, Mir30 shDcc. Each respective control is represented by a dashed red line in the distribution plot. (C) Chart representing the average variation across replicates in the endpoint nuclear displacement on the x-axis from an even probability of 50%:50%, with negative values representing an attraction to the source of the Ntn1 gradient (in red when statistically significant) and positive values representing repulsion (in blue when statistically significant). Unsorted dissociated CGNs were nucleofected and plated on laminin-coated channel microslides for 24 h, then Ntn1 was added unilaterally into the channel 10 min before the start of nuclear tracking for 2 h. Cells were nucleofected with H2B-mCherry, GPI-pHluorin, and the following: LacZ (control), Pard3, JamC, Mir30 shLuc (control), Mir30 shPard3, and Mir30 shJamC. Abbreviations: EGL, external granule layer; ML, molecular layer; IGL, internal granule layer. In (A) through (C), error bars represent the SEM. Statistics: . p ≤ 0.1, *p ≤ 0.05, **p ≤ 0.01, ***p ≤ 0.005, as assessed by an ANOVA followed by a Dunnett post hoc test against the respective controls in (A) and (B) and by a chi-square test in (C) against an even probability of 50%:50%. See also Table S1.

To test the hypothesis that Dcc availability was regulated by the expression of Siah2 in GNPs or Pard3 and JamC in differentiated CGNs, we used Dcc-pHluorin as a reporter for Dcc at the membrane surface and measured the clustering of Dcc in response to Ntn1 being added to the culture medium (Gopal et al., 2016; Matsumoto and Nagashima, 2010). The ratio of the segmented clustered area to the total cell area was calculated over time before and after the addition of Ntn1 (Figure 6B). In the controls cells expressing LacZ, bright Dcc clusters rapidly appeared on the cell surface (Figure 6A, arrowhead), resulting in the clustered area fraction increasing minutes after the addition of Ntn1 (Figure 6C, blue) before slowly returning to normal. Overexpression of Siah2 resulted in a strong decrease in Dcc at the cell surface, whereas both JamC and Pard3 expression were associated with an increased baseline presence of Dcc at the cell surface (Figures 6C and 6D). Under all conditions, adding Ntn1 increased the clustered Dcc membrane fraction and with Pard3 levels being significantly higher than in the controls (Figure 6D). Knockdown of Siah2 resulted in a slight increase in Dcc at the membrane at rest, although this was not significant, and the increased clustered area after Ntn1 addition appeared to decay faster than in the control situation (Figures 6E–6G). Knockdown of both Pard3 and JamC strongly reduced the amount of Dcc presented at the cell surface and the amount that was triggered after the addition of Ntn1 (Figures 6E–6G). These results indicate that the amount of Dcc presented at the cell surface is influenced by the cell polarity pathway and the JamC adhesion molecule.

**Figure 6.**
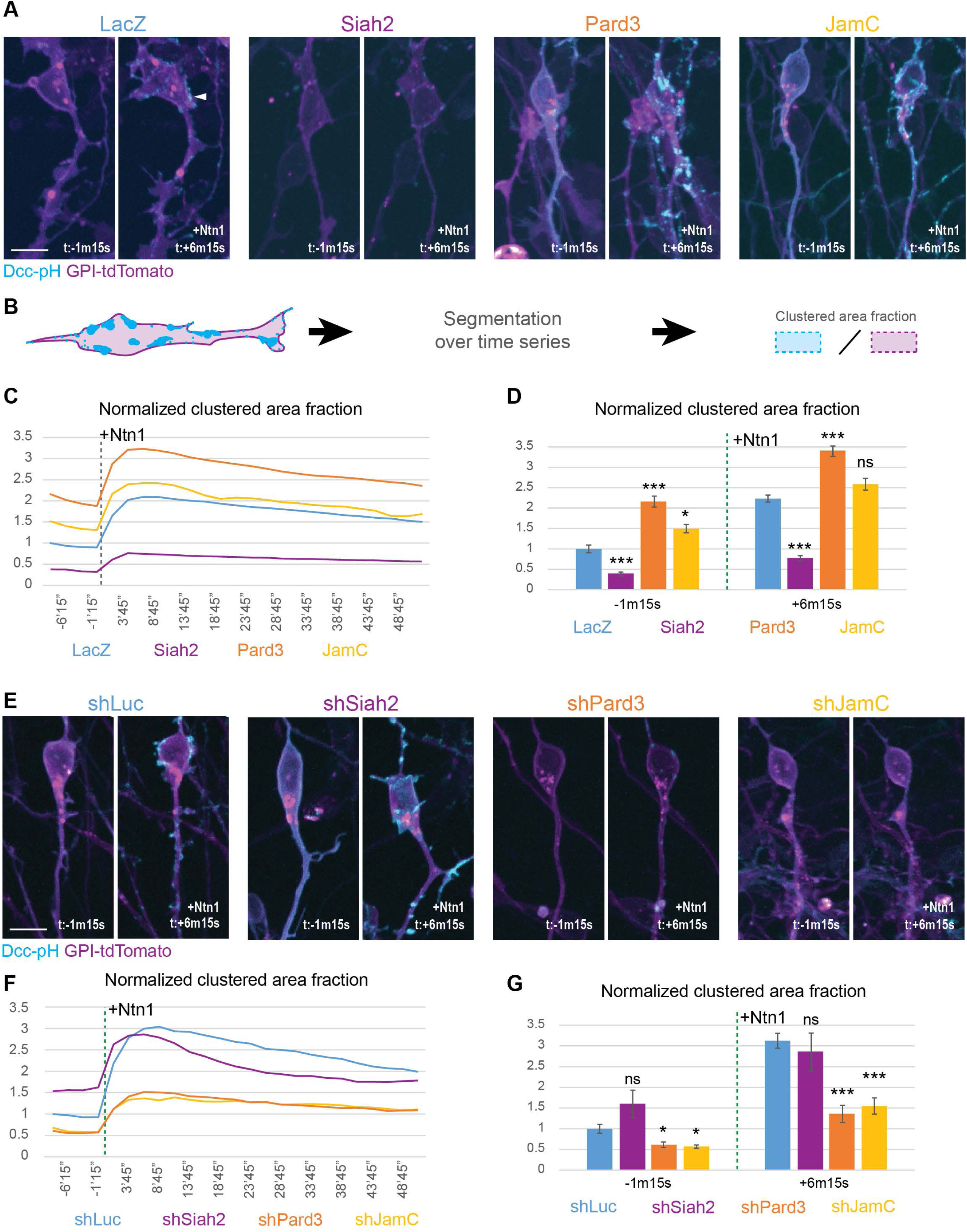
(A) and (E) Spinning-disk confocal live-cell imaging of dissociated granule neurons plated on laminin for 24 h after nucleofection with Dcc-pHlurorin (Dcc-pH) (cyan), GPI-TdTomato (magenta), and one of the following: in (A), LacZ (Ctrl), Siah2, Pard3 or JamC; in (E), Mir30 shLuc, Mir30 shSiah2, Mir30 shPard3, or Mir30 shJamC. Cells were tracked for a total of 1 h at 150-s intervals. Representative pictures for each condition show a maximum projection before (t: −1 m 15 s) and after (t: + 6 m 15 s) the addition of Ntn1 at 200 ng/mL. (B) Schematic representing the segmentation process for the analysis of the time-lapse images in (A) and (E) The Dcc-pH and GPI-tdTomato channels were segmented using Pixel classification with Ilastik. The resulting “clustered area fraction” is the ratio of the area of the segmented Dcc-pH regions (bright Dcc clusters) to the area of the membrane in each field of view for each time point. (C) and (F) Graphs representing the Dcc-pH area fraction over the membrane area, normalized to their respective controls. The different experimental conditions are the same as those in (A) and (E). A dashed line marks the addition of 200 ng/mL of Ntn1 at t0. (D) and (G) Bar charts highlighting data presented in (C) and (F) for a time point before the addition of Ntn1 (t = −1 m 15 s) and for another time point shortly thereafter (t = +6 m 15 s). In (D) and (G), error bars represent the SEM. Statistics: ns, non-significant, *p ≤ 0.05, **p ≤ 0.01, ***p ≤ 0.005, as assessed by an ANOVA followed by a Games–Howell post hoc test against the respective controls. See also Table S1.

## Discussion

In the developing nervous system, progenitor GZ occupancy and subsequent postmitotic neuron migration are considered transitory and separate neuronal maturation phases; this has led to a narrow understanding of how cells within a given lineage at each developmental stage coordinate the GZ exit status of cells whose maturation status varies across the lineage. Whereas GZ exit and radial migration clearly occur at the transition from progenitor proliferation to neuronal terminal differentiation, it remains unknown how “station keeping” between cells in the two states is communicated and interpreted at the cell biological level. For example, increased cell adhesion can sort cells between layers in a developing nervous system tissue by increased cell adhesion molecule expression; however, this is a deterministic event that requires little or no communication between the cells in each layer. In this study, we used the GZ exit of differentiating CGNs of the cerebellum as a model to study how extracellular guidance cue receptors, cellular polarity complexes, and adhesion proteins can intersect, creating a coincidence detection circuit that modulates cell migration behaviors.

We found that Ntn1 signaling affects the somal translocation direction of CGNs. Interestingly, we observed a gradual shift in Ntn1-dependent migratory behaviors; GNPs are unresponsive to this signal, which ultimately transitions to one causing CGNs to be repulsed by a source of Ntn1 as these cells terminally differentiate and mature. Although Ntn1 has previously been described as having a repulsive effect on the migratory behavior of postmitotic neurons (Hamasaki et al., 2001; Yamagishi et al., 2020), this is the first time that a transitional behavior has been described for the same lineage based on the differentiation status of the cells and a change of substrate. It is critical to note that GNPs themselves produce the Ntn1 protein that is repulsive to the CGN progeny of this progenitor population. Therefore, the phenomenon of GNPs ignoring the Ntn1 signal emanating from the GZ itself while CGNs respond to the signal gives rise to the critical station-keeping event that separates these cells from each other in the inner and outer EGL. It is interesting that most neuronal GZ cells express Ntn1 during their peak of proliferation, similar to what is seen in the ventricular zone of the spinal cord and the cerebral cortex (Yamagishi *et al*., 2020; Yung et al., 2015). This suggests that the acquisition of Ntn1 sensitivity by newborn neurons as they differentiate could be a common mechanism to prevent the cells from migrating or projecting back into other GZs (Hamasaki *et al*., 2001; Varadarajan and Butler, 2017).

How is Ntn1 sensitivity ensured during CGN differentiation? We uncovered a coincidence detection circuit whereby JamC and Pard3 are required for Ntn1-induced GZ exit because of the ability of these proteins to increase basal levels of Dcc receptor at the plasma membrane and to increase Ntn1-induced Dcc exocytosis. Although many signaling pathways have been characterized as transducing Dcc signaling in response to Ntn1, this new mechanism ultimately controls the level of CGN repulsion via Ntn1 by affecting the number of signal receptors at the cell surface. Previous results from our laboratory have shown that Pard3 causes JamC exocytosis only in differentiated CGNs, in which it is required for GZ exit. In this study, we have expanded on these results to show that Dcc and JamC extracellular domains interact. Moreover, super-resolution Airyscan microscopy revealed that a subset of Dcc receptors localizes to the periphery of JamC adhesion plaques and that Dcc exocytosis after Ntn1 addition occurs near JamC adhesion plaques. Thus, we have provided functional evidence that Dcc and JamC cooperate with Dcc to promote GZ exit, and the effect of adding Dcc at JamC adhesions suggests a coincidence detection mechanism whereby engaged adhesion receptors at the cell surface provide a template for adding more Dcc. The polarity protein Pard3 also positively affects the amount of Dcc at the cell surface. However, it is unclear whether this is the result of direct action on Dcc exocytosis or a secondary effect of Pard3-mediated exocytosis of JamC, which physically interacts with Dcc through their extracellular domains. The connection between Dcc and the Pard3–JamC polarity-dependent adhesion pathway that creates neuronal adhesion during GZ exit is intriguing. Dcc has been noted to create cell polarity events, including effects on axon formation, and directed invasions of cells in development (Adler et al., 2006; Dominici et al., 2017; Varadarajan and Butler, 2017; Wang et al., 2014); however, no connection between Dcc and the core polarity pathways has been described. Cell migrations frequently require oscillatory mechanisms in cytoskeletal organization or receptor recruitment that are stabilized by cell polarity pathways (Hao et al., 2010). The coincidence detection circuit between Pard complexes and JamC adhesions promotes such oscillatory stabilization: In unmanipulated CGNs, Dcc receptors are quickly cleared from the cell surface after bath application of Ntn1, whereas the clearance of Dcc receptors is enhanced by Pard3/JamC loss of function and retarded by Pard3/JamC gain of function. Stabilization of Dcc receptors at the cell surface clearly creates CGN GZ repulsion, which is essentially a directionally polarized behavior. It will be of great interest to understand how this GZ circuit is precisely controlled at the subcellular level: Although the CGN soma migrates in an Ntn1/Dcc-dependent fashion towards the IGL, CGN parallel fiber axons and their growth cones maintain their position in the molecular layer even in the presence of Ntn1.

Control of Dcc levels and access to the membrane is key to regulating the exit of CGN soma from the EGL. Here, we have confirmed that Dcc is a Siah2 target and have shown that Ntn1-dependent GZ exit can be restrained in GNPs through targeted proteasomal degradation of Dcc by Siah2. Interestingly, Siah2 frequently targets multiple components of a particular cell biological process for degradation (Ong and Solecki, 2017). This appears to be the case for the Ntn1–Dcc GZ exit pathway, as Siah2 also targets Pard3 for degradation, which positively regulates Dcc membrane occupancy. Siah2 expression in GNPs can restrain Ntn1-induced GZ exit by directly targeting Dcc or factors that promote Dcc membrane occupancy. By equally promoting Pard3 degradation and, therefore, inhibiting JamC access to the cell surface (Famulski *et al*., 2010), Siah2 acts as a master regulator of Ntn1 sensitivity in GNPs. It will be interesting to determine whether Siah2 has further roles in modulating Dcc, as previous studies have shown that Ntn1 can promote Dcc ubiquitination (Kim et al., 2005). Indeed, given the rapid disappearance of surface Dcc after Ntn1 addition, Siah2 activity may be regulated by Ntn1 activity or Siah2 affinity for Dcc may be modified as a consequence of Ntn1 binding. It is worth noting that another ubiquitin ligase, Trim9, also regulates Dcc clustering. Trim9 loss of function compromises Dcc clustering, but the loss of clustering is related to unrestrained exocytosis of a host of cargos controlled by soluble N-ethylmaleimide attachment protein receptor (SNARE) proteins due to focal adhesion kinase activation (Plooster et al., 2017). Interestingly, Siah2 degrades Trim9, it may be interesting to dissect whether this additional ubiquitin ligase is part of a larger ubiquitin ligase circuit regulating Dcc signaling in cooperation with Siah2.

### Limitations of the study

shRNA silencing of protein expression is never complete, and our manipulations should be considered a hypomorphic reduction in function.

## STAR★Methods (Cf STAR table)

### Key resources table

### Resource availability

#### Lead contact

Further information and requests for resources and reagents should be directed to and will be fulfilled by the lead contact, David Solecki.

#### Materials availability

All unique/stable reagents generated in this study are available on request from the lead contact.

### Experimental model and subject details

#### Mice

All mouse lines were maintained in standard conditions (e.g., pathogen-free and with continuous access to food/water) in accordance with guidelines established and approved by the Institutional Animal Care and Use Committee at St. Jude Children’s Research Hospital (Protocol no. 483). B6.129(SJL)-Ntn1tm1.1Tek/J, Tg(Atoh1-cre/Esr1*)14Fsh/J, B6.Cg-Gt(ROSA)26Sortm9(CAG-tdTomato)Hze/J, B6.129S-Atoh1tm4.1Hzo/J, and C57BL/6J mouse strains were obtained from The Jackson Laboratory. Neonates were collected on postnatal days 7 through 9 for the studies detailed in the Methods section below or as indicated in the experimental details in the figures. Male and female mice were mixed for the described experiments as no effect of sex on the timing of GZ exit or migration initiation has been observed so far.

#### Method Details

Blinding was not used in data collection.

#### Plasmid vectors

Expression plasmids for LacZ, Pard3a, Siah2, Siah2ΔRING, and H2B-mCherry were subcloned as previously described (Famulski *et al*., 2010; Singh *et al*., 2016) pCIG2 Dcc-pHluorin (Dcc-pH), with a pH-sensitive form of green fluorescent protein (GFP) (Miesenbock et al., 1998) inserted in position 1090, was obtained from Franck Polleux at Columbia University. All new cDNA constructs encoded mouse (*mus musculus*) proteins and were cloned in the laboratory by the overlapping PCR method. See the Key Resource Table for a complete recombinant DNA list.

#### Western blot analysis

HEK293T cells were lipofected (using Lipofectamin 2000; Thermo Fisher Scientific) 1 day before being harvested. Cells were lysed using Lysis/Binding/Wash Buffer (Cell Signaling Technologies) with Halt™ Protease and Phosphatase Inhibitor Cocktail (Thermo Fisher Scientific) and reduced using Laemmli buffer (Sigma-Aldrich, cat. no. S3401). Samples were subjected to polyacrylamide gel electrophoresis using the Bolt™ system (Thermo Fisher Scientific) and transferred to PVDF membranes (Immobilon®-FL PVDF Membrane), blocked in Intercept blocking Buffer, and immunoblotted with appropriate antibodies (see the Key Resource Table for dilutions). For immunoprecipitation and co-immunoprecipitation experiments, the Protein G Immunoprecipitation Kit was used with 1 µg of anti-GFP antibody (RRID: AB_221569) or control IgG (RRID: AB_2722735) in accordance with the manufacturer’s protocol. Samples were then blotted as previously described. HEK293T cells used for blot analysis of the K48-linked ubiquitin signal were treated for 6 h with the proteasome and calpain inhibitor MG132 (50 µM) before being subjected to lysis to see the ubiquitin mark.

#### Ex vivo cerebellar electroporation, organotypic slice culture, and imaging

Cerebella of P7 WT (C57BL/6J) mice were dissected, and the superficial layer of the meninges was removed. Cerebella were soaked in endotoxin-free plasmid DNA suspended in Hank’s balanced salt solution (1–3 μg/μL of each DNA was generally used, with pCIG2-mCherryH2B being electroporated as a nuclear marker for migrating CGNs), transferred to a CUY520-P5 platinum-block Petri dish electrode (Protech International), and electroporated with a CUY21EDIT square-wave electroporator (90 V, 5 pulses, 50-ms pulse, 500-ms interval) (Protech International). Electroporated cerebella were embedded in 4% low-melting-point agarose, and 300-μm sagittal cerebellar slices were prepared using a VT1200 Vibratome (Leica Microsystems). Slices were transferred to Millicell tissue culture inserts (Millipore) and incubated in serum-free medium (FluoroBrite™ DMEM supplemented with 2 mM L-glutamine, 50 cU/mL penicillin–streptomycin, and 1 × B27 and 1 × N2 supplements [Gibco]).

To measure the migration distance of CGNs, cerebellar slices were fixed with 4% paraformaldehyde, mounted on slides using ProLong Gold mountant (Invitrogen), and imaged at 20× with a spinning-disk confocal microscope. Measurements were made with the Amira software, using a self-written script for detecting local maxima of H2B-mCherry–positive nuclei and a Euclidian transform giving the distance from the cerebellar pial surface. Statistical analysis was performed and graphs were prepared with RStudio software.

#### Preparation and nucleofection of CGNs

Briefly, cerebella were dissected from the brains of P7 WT (C57BL/6J) mice, then the tissue was coarsely chopped and treated with a Neural Tissue Dissociation Kit (Miltenyi Biotec). The suspension was layered onto a discontinuous Percoll gradient (35% and 60%) and separated by centrifugation. Cells at the 35%–60% interface were isolated. The resulting cultures routinely contained >95% CGNs and <5% glia. Plasmid expression vectors encoding proteins of interest were introduced into granule neurons via Amaxa nucleofection, using an Amaxa Mouse Neuron Nucleofector Kit in accordance with the manufacturer’s instructions and program O-005. The concentration of pCIG2 expression vectors used for each construct (5 to 25 µg of DNA per 6 million cells) was determined empirically but was consistent across replicates. After cells had been allowed to recover from the nucleofection for 10 min, they were plated in appropriate culture chambers and maintained in culture for 24 h in serum-free medium. For analysis of the Dcc clustering response to Ntn1 (Figure 6), immunocytochemistry (Figures S2B and S4B), and live imaging (Figures 4A–4C; Figures S4C and S4D), we used 6-cm dishes with glass bottoms that had been treated with poly-L-ornithine then coated with laminin at 1 µg/cm^2^.

#### Atoh1-eGFP–sorted cells

Mouse pups homozygous for Atoh1::GFP (B6.129S-Atoh1tm4.1Hzo/J; The Jackson Laboratory, JAX: 013593) were collected at P7, and cerebellar granule neurons were isolated after dissociation by the methods described above. “High” and “low” populations were sorted, based on their GFP intensity, with a cell sorter (FACSAria Fusion, BD Biosciences; 85-μm nozzle, 35 PSI). The GFP intensity distribution profile was constant (Figure S2A), and the gating criteria were consistent across replicates.

#### Migration assays on channel microslides

For migration assays, microslides (µ-Slide I; Ibidi, cat. no. 80106) were treated with poly-L-ornithine and then coated with laminin at 1 µg/cm^2^ or vitronectin at 2 µg/cm^2^. Two million nucleofected CGNs or freshly sorted “low” or “high” Atoh1::GFP cells (prepared as described above) were then added to the channel and left to incubate. After 24 h, before the start of the time-lapse image acquisition, the medium was replaced twice by aspirating the existing medium from one end of the track and adding 100 µL of warm, fresh medium at the other end. After a 10 min incubation, the same procedure was repeated, but this time adding only 50 µL of serum-free medium containing Ntn1 to a final concentration of 250 ng/mL or the equivalent volume of 1× PBS as a control. Cells were then tracked every 150 s for 130 min after Ntn1 addition in a spinning-disk confocal microscope with environmental control. Cell body migration was tracked manually with SlideBook (Intelligent Imaging Innovations) by using the H2B-mcherry signal (in nucleofected cells) or differential interference contrast (DIC) (in Atoh1-sorted cells). Cells showing less than 10 µm of total displacement over the time-lapse period were analyzed.

#### Cerebellar immunohistochemistry

Postnatal brains were fixed by immersion in 4% paraformaldehyde at 4°C overnight, washed five times in 1× PBS for 24 h, then cryoprotected in PBS containing 30% sucrose. Histologic sagittal sections were cut on a cryostat and pre-blocked for 1 h in PBS with 0.3 M glycine, 0.1% Triton X-100, and 10% normal donkey serum. Sections were incubated overnight at 4°C with the primary antibodies (see the Key Resource Table for dilutions). This was followed by incubation at room temperature for 1 h with the appropriate Alexa Fluor–labeled secondary antibody (Invitrogen; diluted 1:1000) before mounting. Images were acquired with a spinning-disk microscope, using SlideBook software.

#### Pulse-chase assay in Atoh1::CreERT2; Ntn1^flox/flox^

Tamoxifen in corn oil was injected intraperitoneally (at 100 mg/kg body weight) at P0, P1, and P2. For pulse-chase migration assays, mice were injected intraperitoneally with 50 mg/kg of EdU 48 h before tissue collection. EdU incorporation was assayed with the Click-iT assay (Invitrogen) according to the manufacturer’s instructions.

#### Quantification and statistical analysis

Standard image processing and analysis were carried out with Amira (Thermo Fisher), SlideBook, or Fiji software. For cluster area analysis, pixels were classified with a random forest classifier, using Ilastik for sum projection after drift and Fiji for photobleach correction. Analyzed metrics of quantitated data are expressed as the mean ± SEM or as an adjusted ratio of clustering relative to the control average at the initial time point. The Student *t*-test was used to compare two groups. A one-way analysis of variance (ANOVA) was used for multiple comparisons with the Dunnett post hoc test against controls (for ex vivo slices) or with the Games–Howell post hoc test (for cluster area analysis). A non-parametric post hoc test was used for the cluster area analysis because the data did not satisfy the homogeneity of variance. The chi-square test was used in the channel microslide migration assay to test the results against an even probability (0.5) of the relative endpoint displacement on the axis of the length of the channel being either toward or away from the source of the gradient. In this assay, data were expressed graphically as the percentage variation from 50%, showing the mean ± SEM across replicates. All statistical test assumptions were verified when required by the test.

#### Data and Code Availability

Scripting was done with Fiji and Amira. Original imaging data is available upon request.

## Acknowledgments

We thank Franck Polleux for the original Dcc-pHluorin construct and Luke Lavis for sharing the Lampshade Magenta dye. Zeiss 980 Airy Scan images were acquired at the Cell & Tissue Imaging Center (CTIC), which is supported by SJCRH and NCI P30 CA021765. Drs. Aaron Taylor, Aaron Pitre, and George Campbell provided consultation and support for imaging experiments in CTIC. Keith A. Laycock, Ph.D., ELS, did the scientific editing of the manuscript. The Solecki laboratory is funded by the American Lebanese Syrian Associated Charities (ALSAC) and by grants 1R01NS066936 and R01NS104029 from the National Institute of Neurological Disorders (NINDS). The content of this manuscript is solely the responsibility of the authors and does not necessarily represent the official views of the NIH.

## Author Contributions

C.L designed and carried out all the experiments and analyses. L.S. tracked Atoh1 sorted CGNs and participated in the analysis of Netrin1 in vivo experiments. D.R.S. carried out pilot imaging experiments examining the configuration of surface recruited DCC. T.L.L. constructed the DCC pHluorin construct. N.T. maintained mouse colonies used in all experiments. D.H. prepared CGNs and reagents for pilot experiments. D.J.S. carried out pilot experiments and conceived and supervised the study. All authors drafted or edited the manuscript.

## Graphical abstract

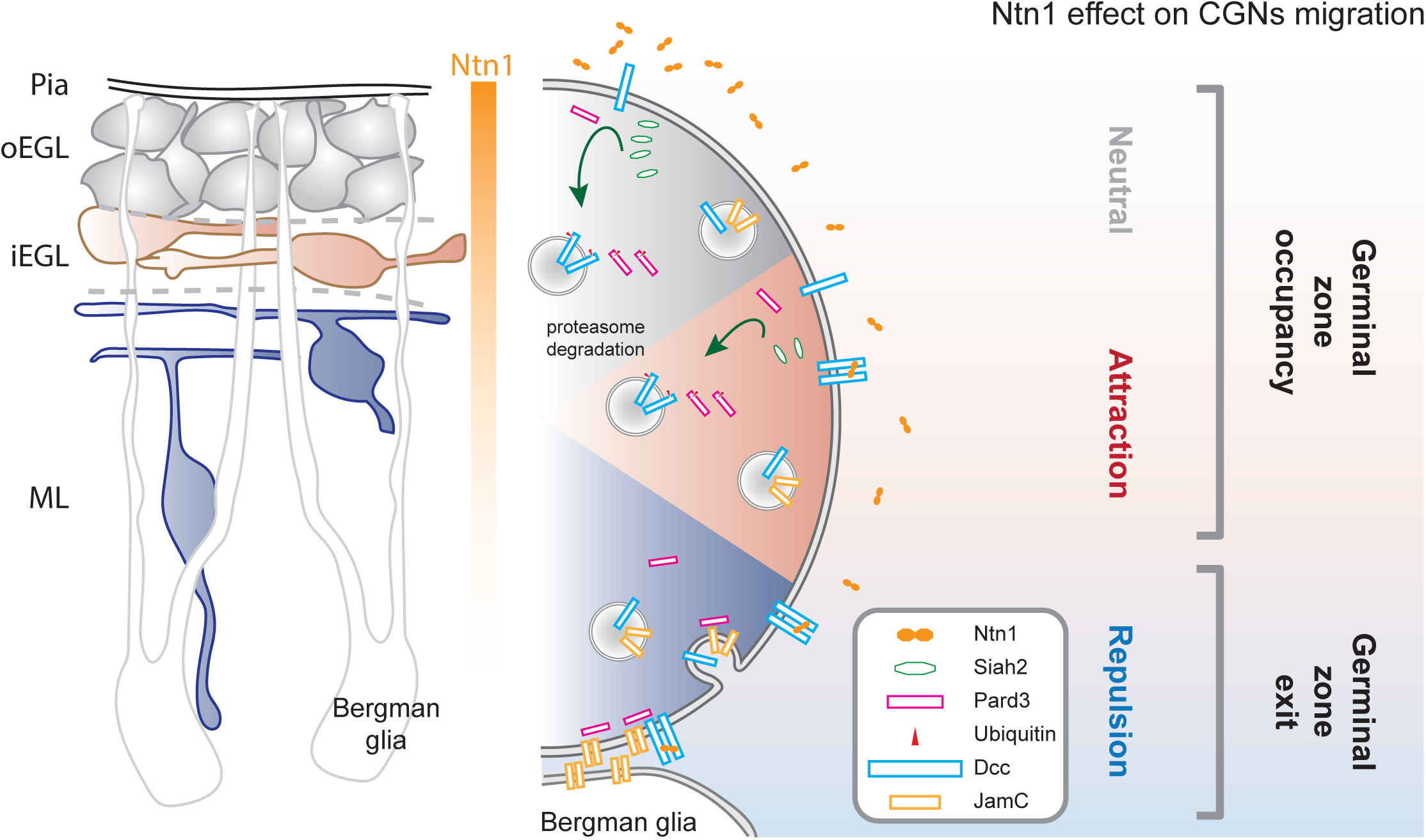

---

**Figure S1.**
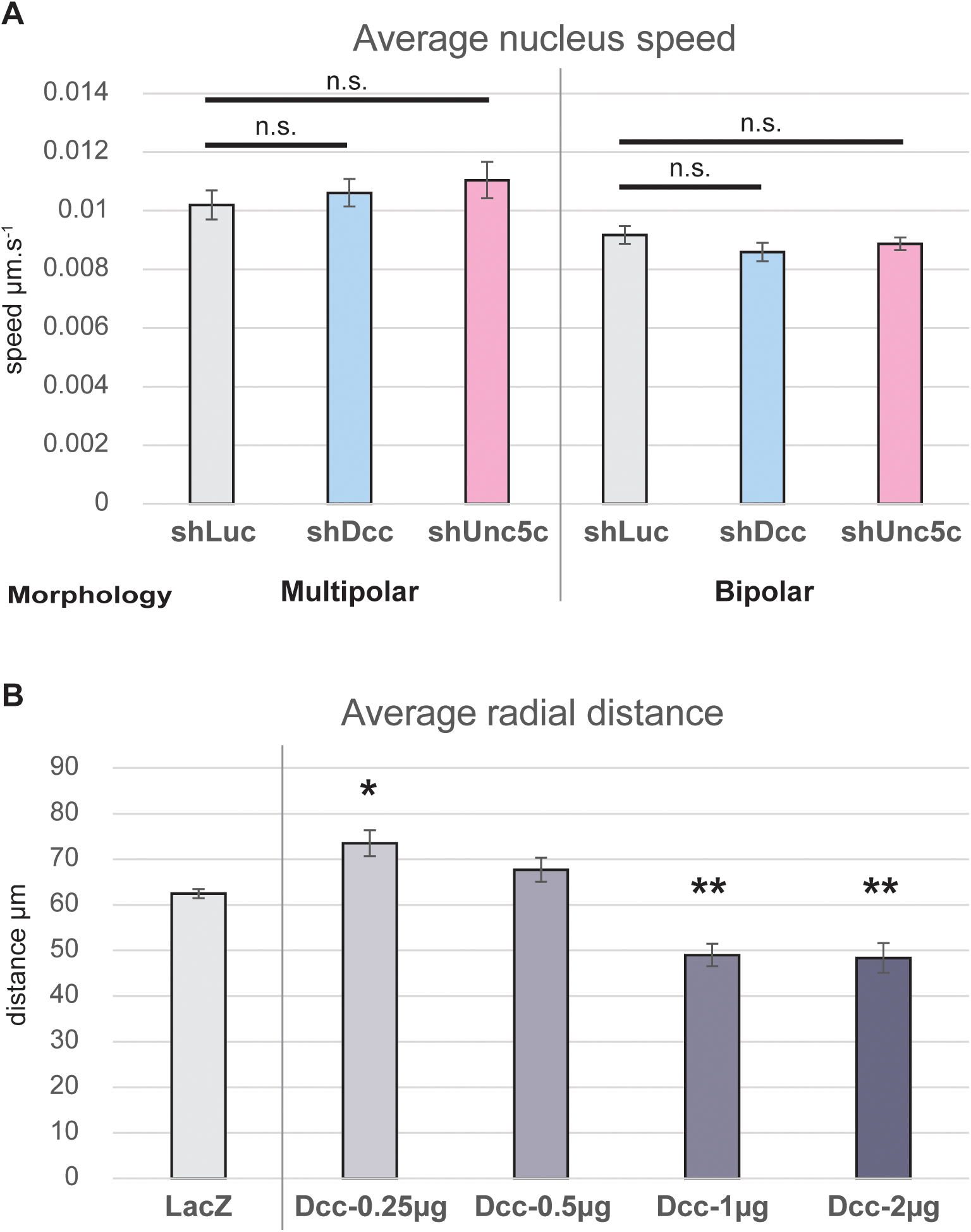
(A) Bar chart representing the average speed of the nuclei (in µm.s^−1^) in dissociated cerebellar granule neurons plated on laminin for 24 h and tracked over 2 h at 150-s intervals. Cells were nucleofected with Centrin2-venus, H2B-mCherry, and Mir30 shRNA against luciferase (Luc, gray), Dcc (blue), or Unc5c (pink), and the results were separated based on cellular morphology. Cells with a progenitor-like morphology without clear polarity or a long neurite were categorized as multipolar, and cells with a defined polarity and a neurite longer than 30 µm were categorized as bipolar. (B) Bar chart representing the average radial distance of nuclei of P7 mouse cerebellar cells electroporated in an *ex vivo* cerebellar slice assay, after 48 h in culture. The results show the effect of increasing concentrations of Dcc on germinal zone exit. Total DNA concentrations were adjusted to match the highest concentration of Dcc with the corresponding amount of LacZ. In (A) and (B), error bars represent the SEM. Statistics: n.s., non-significant, *p ≤ 0.05, **p ≤ 0.01, as assessed by a Student *t*-test in (A) and an ANOVA followed by a Dunnett post hoc test in (B). See also Table S1.

**Figure S2.**
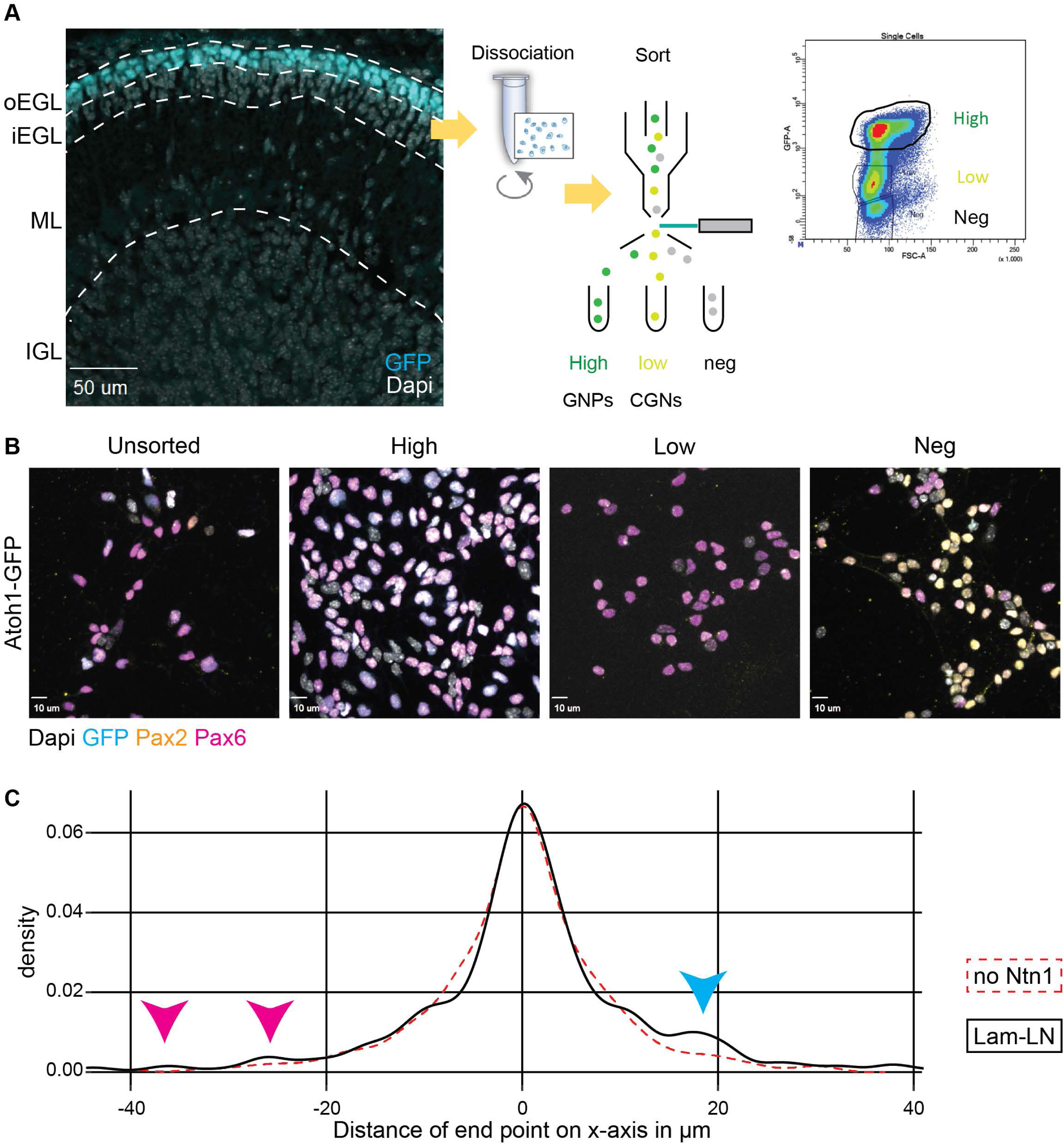
(A) (Left) Immunohistochemical staining with an antibody against GFP in sagittal cryo-sections of Atoh1-GFP mouse cerebellum at P7, showing the restricted expression of Atoh1-GFP protein in the oEGL. (Right) Schematic representing the protocol used to isolate and sort the different cell populations from Atoh1-GFP mice cerebella. The GFP intensity profile shows three clusters, high, low, and negative (Neg), and the cells were separated accordingly. (B) Immunocytochemical staining for Pax2, Pax6, and GFP in the different sorted Atoh1-GFP populations from (A). (C) Density profile of the x-value for endpoint nuclear migration on channel microslides coated with laminin in response to an Ntn1 gradient for the LN population (black line) and the no-Ntn1 controls (red dashed line) (see Figure 2E). Arrowheads indicate cells whose migratory behavior showed strong attraction (magenta) or repulsion (cyan) in response to Ntn1. Scale bars in (A) and (B) represent 50 µm and 10 µm, respectively. Abbreviations: oEGL, outer external granule layer; iEGL, inner external granule layer; EGL, external granule layer; ML, molecular layer; IGL, internal granule layer; GNPs, granule neuron progenitors; CGNs, cerebellar granule neurons; Neg, negative.

**Figure S3.**
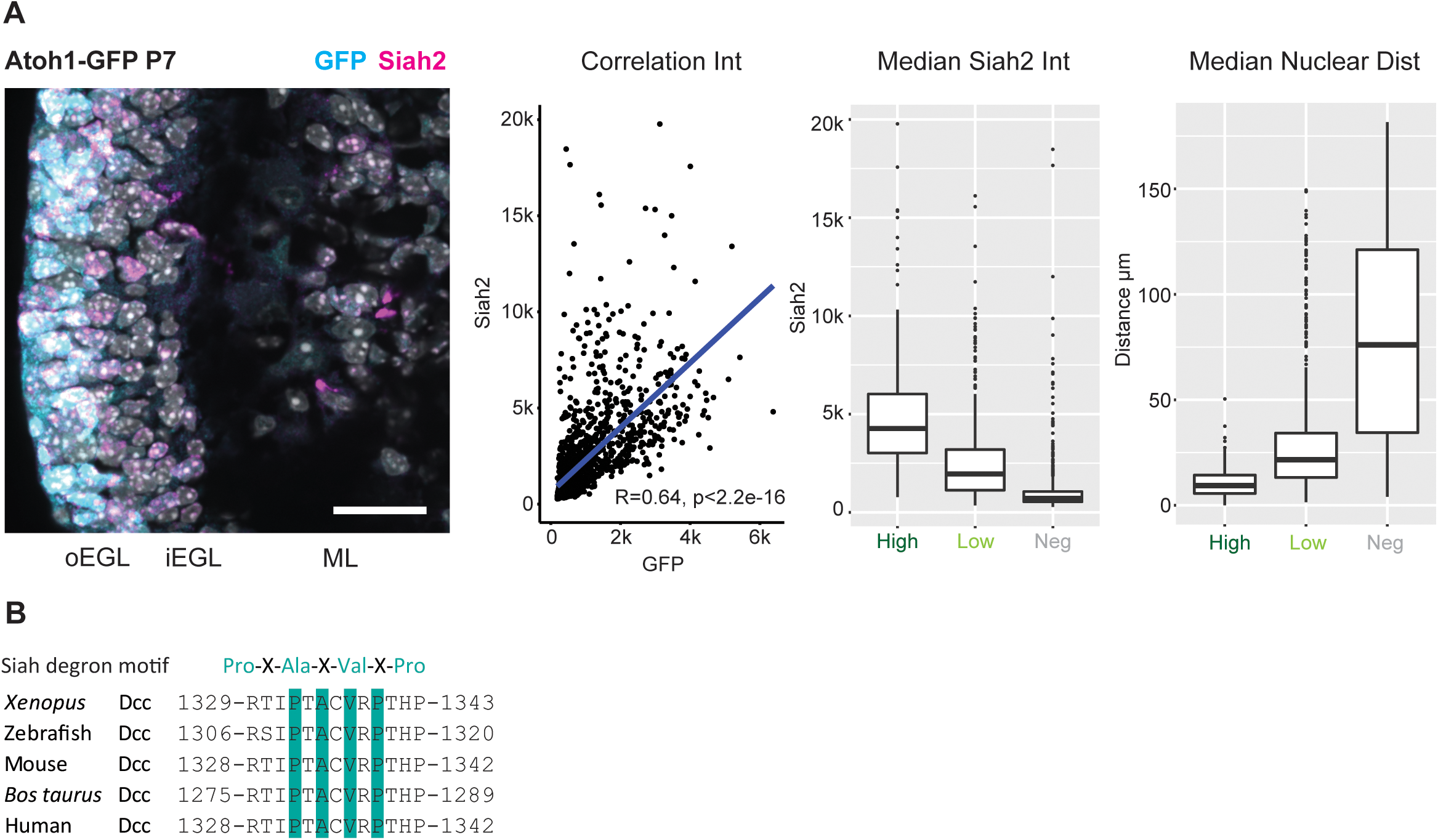
(A) (Left) Immunohistochemistry staining of sagittal cryo-sections of Atoh1-GFP mouse cerebellum at P7, against GFP (Cyan) and Siah2 (Magenta), counterstained with Dapi (Gray). First Graph shows the significative correlation between GFP and Siah2 signal intensity in segmented cell bodies. Second and Third graphs shows the boxplot distribution of Siah2 intensity (second) and the distance from the cerebellar surface (third) for the 3 populations of granule neurons based on GFP intensity: “High”, “Low” and “Neg” (negative). (B) Sequence alignment of different Dcc orthologues, showing the conservation of the Siah degron motif, Pro-X-Ala-X-Val-X-Pro, within the intracellular P2 domain of Dcc.

**Figure S4.**
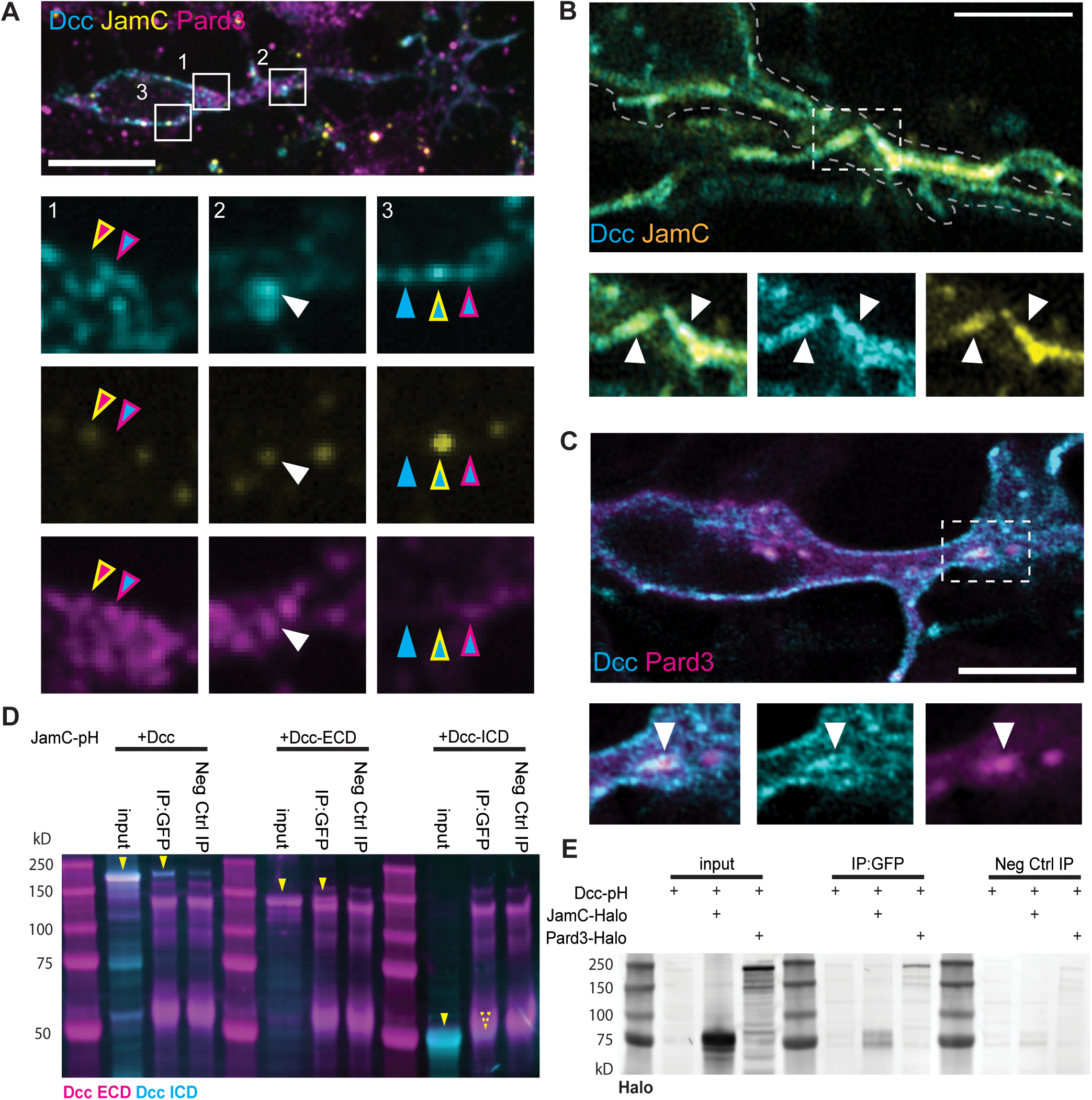
(A) Confocal imaging and single focal plane images of a dissociated cerebellar granule neuron plated on laminin and maintained in culture for 24 h before fixation and immunohistochemical staining with antibodies against Dcc (cyan), JamC (yellow), and Pard3 (magenta). Magnifications of frames 1–3 are shown below the top image, with arrowheads highlighting areas where staining for Dcc and Pard3 (cyan and magenta arrowhead), Dcc and JamC (cyan and yellow arrowhead), Pard3 and JamC (yellow and magenta arrowhead), and Dcc, JamC, and Pard3 (white arrowhead) overlaps, or where staining for Dcc does not overlap with staining for other proteins (cyan arrowhead). (B) and (C) Single focal plane image from Airyscan confocal live imaging of CGNs nucleofected with Dcc-pHluorin (cyan) and JamC-SNAP (yellow) constructs (B) or Dcc-pHluorin (cyan) and Halo-Pard3 (magenta) constructs (C). Phluorin and the SNAP dye used here are both pH sensitive, emitting fluorescence only if exposed to a neutral pH. Thus, they show only proteins currently at the cell membrane surface. The regions contained in the dotted rectangles are magnified below, showing where the Dcc-pH and JamC signal overlap at the membrane surface at a point of contact between two neurites (B, white arrowheads) or where the Dcc-pH and cytoplasmic Pard3 signal overlap in the proximal dilation of a CGN (C, white arrowheads). (D) Western blot of lysates of HEK293T cells after lipofection with JamC-pHluorin (JamC-pH), Dcc, Dcc extracellular and transmembrane domain (Dcc-ECD), and Dcc intracellular domain (Dcc-ICD). Cell lysates were immunoprecipitated with an antibody against GFP (IP: GFP) or a negative-control rabbit IgG (Neg Ctrl IP). The blots were immunostained with two different antibodies against Dcc: one against the extracellular domain and one against the intracellular domain. Yellow arrowheads highlight the expected band sizes with Dcc, Dcc-ECD, and Dcc-ICD. (E) Western blot of lysates of HEK293T cells after lipofection with Dcc-pHluorin (Dcc-pH), JamC-Halo, and Pard3-Halo. Cell lysates were immunoprecipitated with an antibody against GFP or a negative-control rabbit IgG. The blot was immunostained with an antibody against the Halo tag. The blot shows enrichment of JamC-Halo and Pard3-Halo in the eluate when Dcc-pH is pulled down with anti-GFP antibody. Scale bars in (A) represent 10 μm; those in (B) and (C) represent 5 µm.

**Figure S6.**
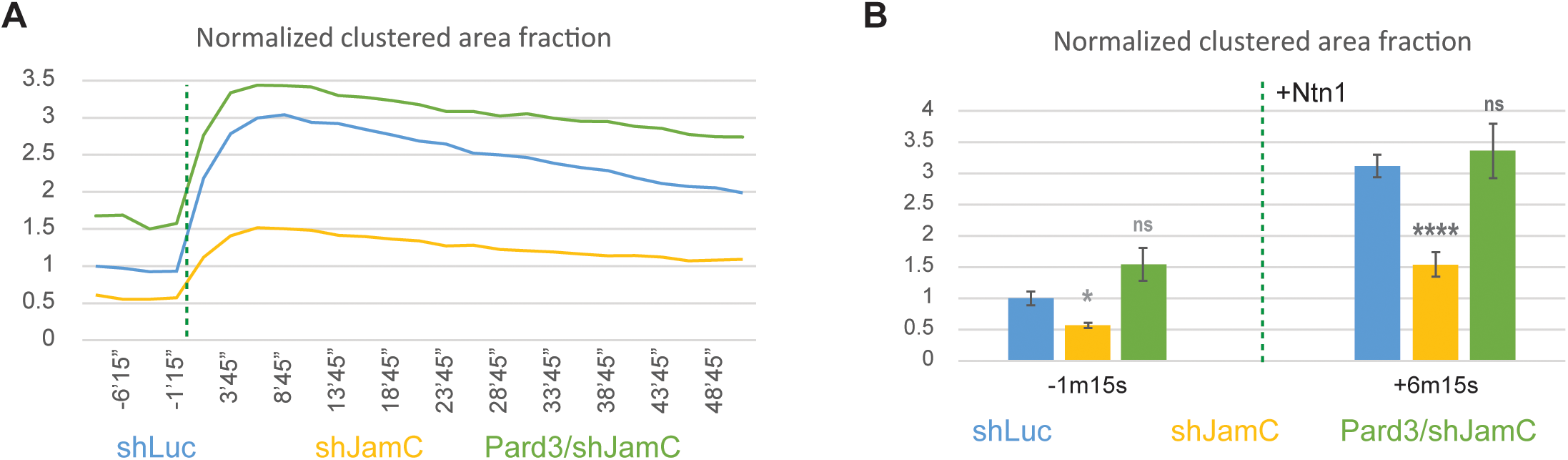
(A) Graph complementing the results of Figures 6E–6G, representing the Dcc-pH area as a fraction of the total membrane area, normalized to the control. Here, there is an additional experimental condition, with CGNs being nucleofected with Pard3 and Mir30 shJamC. The dashed line marks the addition of 200 ng/mL of Ntn1 at t0. (B) Bar chart highlighting data presented in (A) for a time point before the addition of Ntn1 (t = −1 m 15 s) and another one shortly thereafter (t = +6 m 15 s). In (B), error bars represent the SEM. Statistics: ns, non-significant, *p ≤ 0.05, **p ≤ 0.01, ***p ≤ 0.005, as assessed by an ANOVA followed by a Games–Howell post hoc test against the control (shLuc). See also Table S1.

